# Voltage-gated ion channel diversity underlies neuronal excitability and nervous system evolution

**DOI:** 10.1101/2025.02.15.638427

**Authors:** Jose Davila-Velderrain, Lena van Giesen

## Abstract

All nervous systems function via electrical excitability mediated by ion channels. However, channels and excitability are both ancient, preneuronal features that many cells use to rapidly adjust behavior. We systematically studied the evolutionary paths to neuronal excitability by characterizing the voltage-gated ion channel complements (VGL-chanomes) of 623 organisms, dissecting their expression patterns in 11 whole-body cell atlases and 3 entire nervous systems, and recording electrical properties of “primitive” neurons in the sea anemone. We find a disconnect between ion channel availability and organismal or nervous system complexity and find instead an association with lifestyle and behavior. Cell type restricted chanome expression predated the emergence of nervous systems in multicellular organisms. Multiple gene-family expansions and contractions independently specialized or diversified VGL-chanomes leading to a surprising convergent pattern: not the number of channels but their diversity and restrictive recruitment is a hallmark of neuronal complexity. These findings suggest that the evolution of highly complex nervous systems was not a stepwise progression of expanding complexity.

## Introduction

All organisms sense and process information from their internal and external environment. Appropriate real-time behavioral responses often require rapid signaling, especially in large animals. The fastest, millisecond-scale signaling modality used by organisms is electricity. This bioelectricity is generated by semipermeable membranes that separate ions in an electrically polarized configuration that accounts for the negative membrane potential (voltage) of cells^1^. Cells of a special class (excitable cells) are able to actively modify this potential to encode and transmit information by opening and closing different classes of ion channels (**Fig. 1a**)^2–4^. The opening probability of some of these channels (voltage-gated) is itself affected by voltage changes, thus establishing a feedback control mechanism to actively regulate membrane potential^5–7^. The regulated dynamics of membrane (re)depolarization provides organisms with a mechanism for rapid information transmission and processing. Our ability to sense, move, feel, and think relies on this mechanism^8–12^.

**Figure 1.**
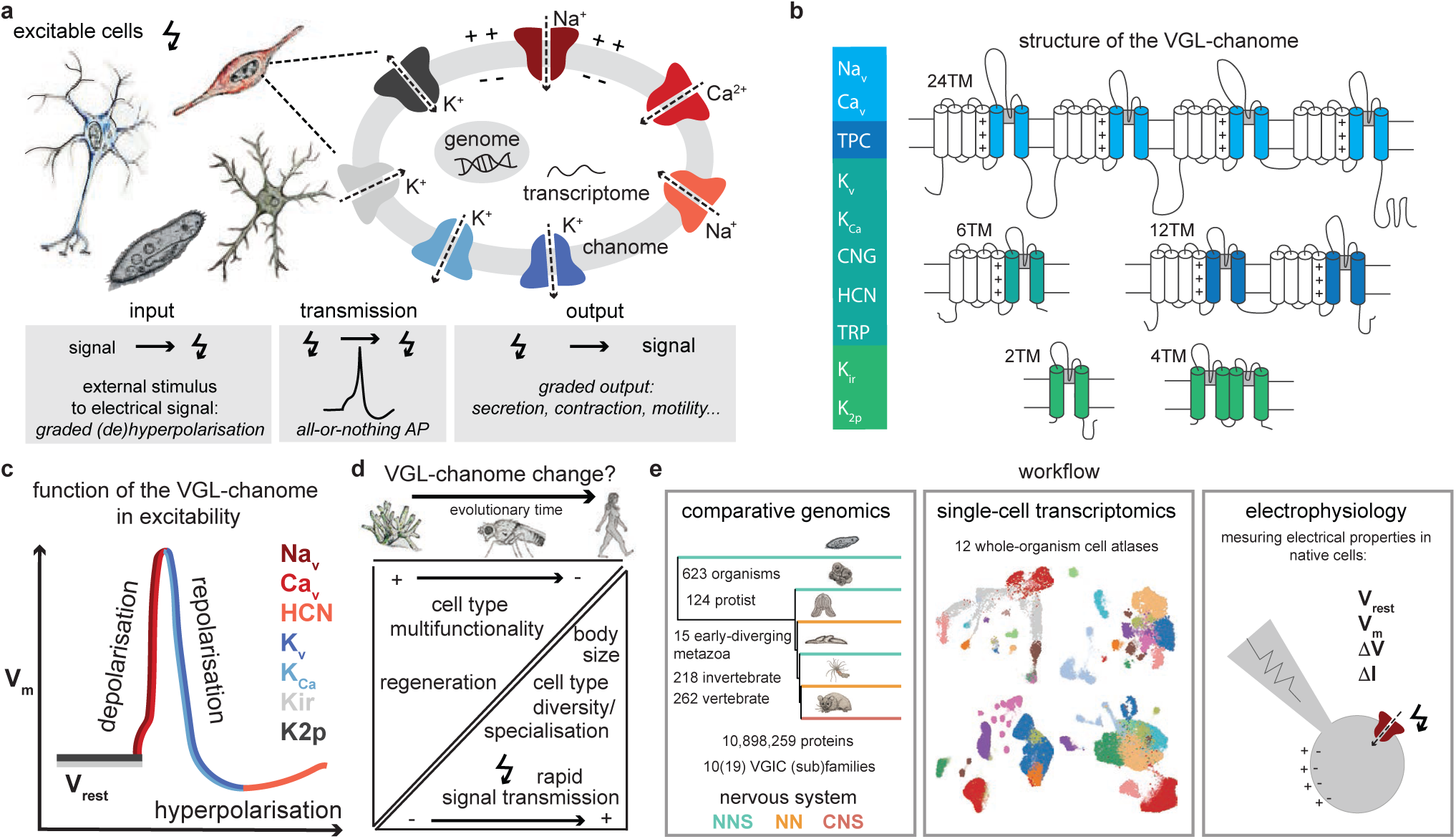
Evolution of excitable cells and voltage-gated ion channels. **a)** Excitable cells contain specific types of voltage-gated ion channels that enable dynamic and regenerative membrane potential changes, enabling the rapid encoding and transmission of signals and subsequent transduction into graded output. **b)** These ion channels belong to the voltage-gated-like (VGL) superfamily that contains related members with distinct membrane topologies. **c)** In excitable cells, specific members of the VGL-chanome are responsible for defined changes in membrane potential (resting membrane potential Vrest; de-, re- or hyperpolarization) that enable dynamic signal coding. **d)** How the VGL-chanome has changed over evolutionary time to accommodate cell type multifunctionality, body size, and regenerative capacities in diverse organisms is unknown. **e)** We systematically analyzed VGL-chanome dynamics in terms of genome content, expression patterns, and physiological function across a variety of metazoans and protists using available genomic and single-cell transcriptomic data.

Electrical excitability is most often studied in the sensory receptor cells, neurons, and muscle cells of organisms with a nervous system^13,14^. However, bioelectric signaling is implicated in many other physiological processes, including the spread of protective and locomotory responses by neuroid (non-nervous, non-muscular) conduction^15^, morphogenetic information processing ^16^, and microbial community coordination ^17,18^. It also underlies the sensory-motor responses of unicellular protozoans. In *Paramecium* voltage-gated ion channels segregate to specific subcellular compartments (e.g., front, back, or cilia) bypassing the need for multicellular specialization^19,20^. Classical electrophysiological experiments have demonstrated functional “nervous” ion channels in protozoa, yeast, and bacteria^21^. Complex behavioral control and bioelectric molecular toolkits thus seem to have predated the emergence of nervous systems or even multicellularity^13^.

Genomic and phylogenetic studies have confirmed these observations. Genomic studies uncovered a surprising diversity of ion channels in different organisms^22,23^. Voltage-gated ion channels (VGICs) form part of a gene superfamily specialized for electrical signaling which also includes VGIC-like (VGL) proteins that are structurally similar but lack voltage-dependent kinetics. All share a modular organization of protein motifs consisting of combinations of 2 or 6 transmembrane (TM) domains, and their total complement in an organism has been referred to as the VGL-chanome (**Fig. 1b**)^24^. It includes the classical voltage-gated Na^+^, K^+^, and Ca^2+^ channels, cyclic-nucleotide gated channels (CNG), non-voltage-gated potassium channels (Kir, K2p), Transient Receptor Potential (TRP) channels, as well as channels in which voltage acts in concert with ligands (K_Ca_ and HCN) -- all encoded by multiple genes in mammals. Seven VGL-channel families are particularly relevant for intrinsic electrical excitability and are largely responsible for dynamic membrane potential regulation via de-, re-, or hyperpolarization: voltage-gated sodium (Na_v_), calcium (Ca_v_), and hyperpolarization-activated cyclic nucleotide-gated (HCN) channels for depolarization; voltage-gated (K_v_) and calcium-activated potassium (K_Ca_) channels for re- and hyperpolarization; and inward-rectifier (Kir) and two-pore domain potassium (K2p) channels for resting membrane potential (**Fig. 1c**). Phylogenetic studies have shown that VGL-chanome members are found in all domains of life^21,25^, and that channels mediating electrical excitability in animals predated neurons^26–28^. Ion channel types relevant for nervous system function have an ancient origin and are thought to have undergone independent gene expansions in animals with diffusive nervous systems (Ctenophores, Cnidarians)^29^. The expansion of specific families may also have coincided with increased brain complexity in mammals^29,30^.

In addition to genome composition, the combinatorial way in which excitable cells use the ion channel toolkits available in their genome is important for understanding their functional specialization. The cell type specificity, expression variability, and co-expression patterns of VGL-chanome proteins largely determine the bioelectrogenic and computational properties of excitable cells^2,31–33^. Because both ion channels and cell types changed drastically in number and complexity during animal evolution, studying the interplay of excitable VGL-chanomes, excitable cells, and nervous system organization is not trivial (**Fig. 1d**). The question of how ion channels expanded and distributed across specialized cell types as organismal structure complexified and nervous systems emerged remains open and is not addressable by comparative genomics and phylogenetics alone. Since VGICs are the molecular basis of electrical excitability in all organisms, the systematic study of VGL-chanome composition across many diverse organisms, and its selective use in specific cell types within representative organisms, provides an opportunity to investigate global evolutionary patterns of electrical excitability and its apparent specialization for nervous system function. We leveraged extensive genomic and transcriptomic resources to revisit these fundamental questions following a systematic integrative approach. We combine comparative genomics, whole-organism single-cell transcriptomics, protein modeling, and electrophysiology to globally characterize the VGL-chanome composition of 623 metazoan and protozoan genomes, their patterns of variability across organisms, and their restrictive expression and function in specialized excitable cells (**Fig. 1e**).

## Results

### Characterizing the VGL-chanome of metazoans and protists

As a first step to investigate the interplay between cellular excitability, cellular specialization, and nervous system evolution, we characterized the genomic repertoire of VGL-channels in 623 organisms. We considered early diverging extant metazoans (*Porifera*, *Ctenophora*, *Placozoa*, *Cnidaria*) and their close unicellular relatives (*Filasterea* and *Choanoflagellatea*) to study compositional changes at the onset of multicellularity and the early evolution of nervous systems (**Fig. 1e**). We included 124 protists to examine the extent to which single-celled eukaryotic organisms without a nervous system are nonetheless equipped with electrically excitable molecules (**Supplementary Table S1)**. We identified the VGL-chanome complement of each genome by performing systematic genome-wide searches for structurally related amino acid sequences using the common pore region of mammalian VGL-channels as a template. Following^24^, we built hidden markov model (HMM) profiles of 18 ion channel subfamilies using sequences corresponding to the minimal pore structure (**Supplementary Table S2**). We interrogated these models against the complete set of annotated proteins of each organism (10,898,259 proteins in total) and filtered out low quality matches based on protein length, predicted membrane topology, matching scores, and nearest-neighbor matching criteria (**Supplementary Fig. S1a,** Methods). This search strategy identified 62,374 VGL-channel candidates across 623 genomes (100.9 VGL-coding genes per genome on average) supported by multiple sources of evidence (**Supplementary Table S3**). HMM profiles accurately recovered well-characterized VGL-chanomes in human, mouse, fly, and worm; with only few low-scoring false positive hits that did not pass subsequent filtering criteria (e.g., 8 out of 153, 94.77% accuracy in human) (**Supplementary Fig. S1b)**. Length, conservation, and membrane topology features of identified candidates are consistent with known VGL-channel structures (**Supplementary Fig. S1c,d)**. Candidates have 875.56 residues on average, highly significant matches with mammalian VGL-chanome members (average E-value < 4.6 x 10^-5^, *phmmer*) and Pfam ion channel domains (average E-value < 3.5 x 10^-5^, *hmmserch* pfam Ion_trans), best-hit matches with mammalian VGL-channels (96.54% of the candidates), and the majority (86.59%) have a predicted topology with 2, 4, 6, 8, 12 or 24 transmembrane (TM) domains (**Supplementary Fig. S1c**). A 6TM topology was the most frequent (45.5%), consistent with potassium channel families being the most common in animal genomes^34^. We predicted the membrane topology of all candidate channels using the deep learning protein language model DeepTMHMM^35^, which accurately recovered the known topology of mammalian VGL-channels directly from sequence (**Supplementary Fig. S1d**). VGL-channel annotations, protein sequences, and topology predictions are reported in **Supplementary material**.

### VGL-chanome size does not reflect organismal complexity

Having many ion channels is commonly interpreted as indicative of the many possible genetic ways of constructing an excitable cell in an organism^32^. We grouped organisms based on VGL-chanome size to investigate its patterns of association with organismal complexity and nervous system organization. VGL-chanomes vary greatly in size, from a single channel in the bacterium *B. subtilis* (included for reference) to a maximum of 385 VGL-channel candidates in the ray-finned fish *S. rhinocerous* (**Fig. 2a**). Organisms also vary greatly in their total number of protein-coding genes (3,000 to 50,000 range). Both chanome and genome size vary within taxa as well, in particular in protists, cnidarians, and bilaterians (**Supplementary Fig. S2a,b**). Therefore, estimates of chanome size based on direct channel gene counts and average or median estimates within taxa might not accurately reflect diversity patterns (**Supplementary Fig. S2c**). To account for this variation, we represent the size of the VGL-chanome of a given organism as the total number of VGL-channels per every thousand protein-coding genes in its genome. We refer to this metric as the CPT score (channel count per thousand genes) and interpret it as a proxy for the genomic investment an organism places in the capacity to produce VGL-channels. The CPT distribution revealed 4 regimes of increasing investment (Gaussian mixture model, Bayesian information criterion BIC -2791.472) (**Fig. 2a**). The genome of organisms in the first, second, third, or fourth group (g1-g4) encodes on average 0.60 (n=85), 3.84 (n=238), 7.53 (n=292), or 10.95 (n=8) VGL- channels per every 1000 protein-coding genes.

**Figure 2.**
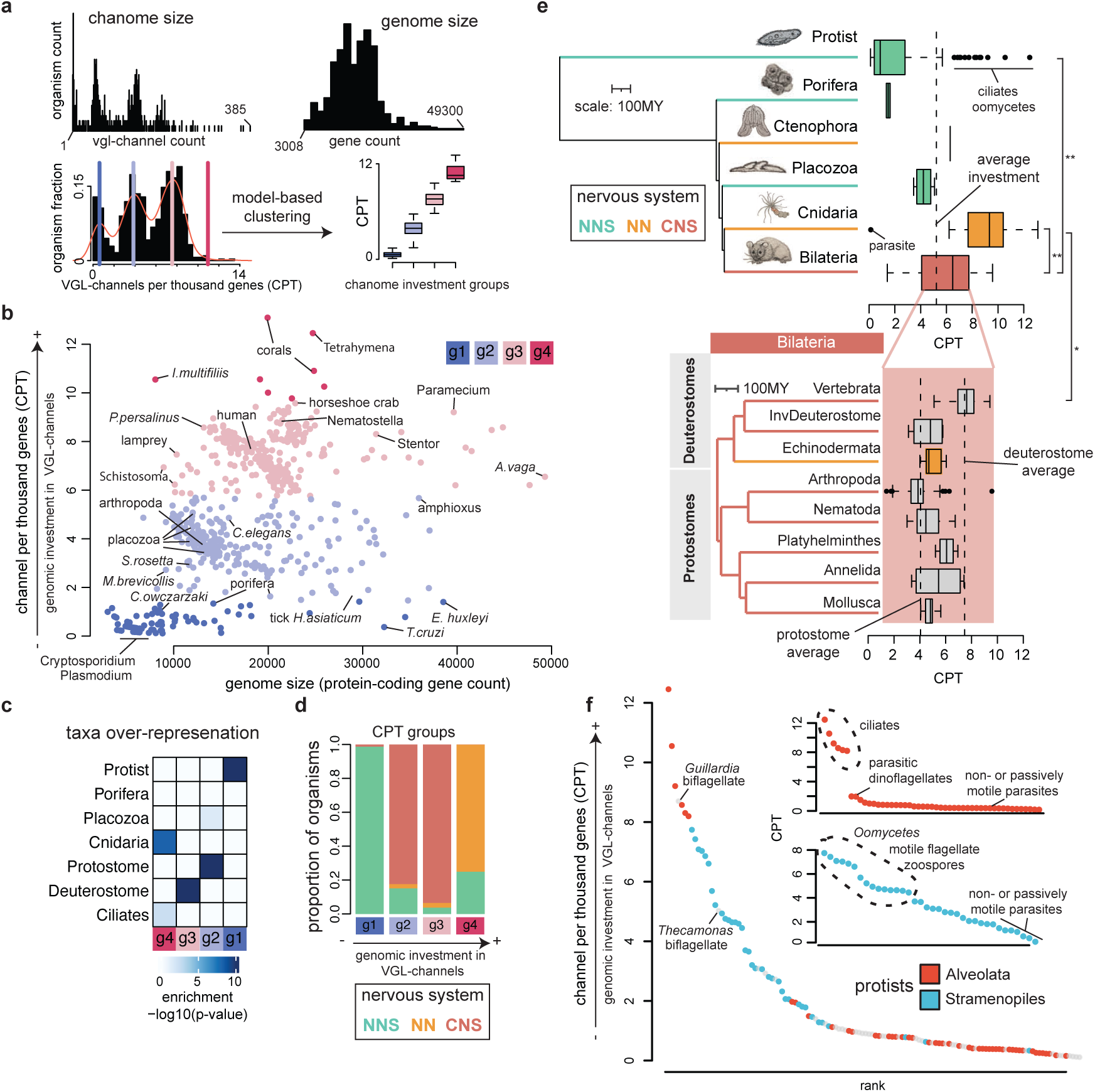
Associations between VGL-chanome size, organismal complexity, and nervous system organization. **a)** Chanome and genome size distributions across 623 organisms. The VGL-channel count per every thousand protein-coding genes in a genome (CPT) is used to estimate and compare effective chanome sizes across organisms. Chanome sizes cluster into 4 chanome investment groups. The CPT score is interpreted as a proxy for the genomic investment an organism places in the capacity to produce VGL-channels. **b)** Relationship between chanome (CPT) and genome size highlighting investment groups (color-coded) and representative organisms. Data points represent CPT values per organism (n=623). **c)** Over-representation of major taxa within investment groups. *P* values were calculated using binomial tests. **d)** Relationship between nervous system organization and CPT investment groups. **e**) Distribution of organismal CPT values across taxa. Data points represent CPT values per organism. *P* values were calculated using Wilcoxon two-sided signed-rank tests. **f)** Rank-ordered CPT values for protists from groups *Alveolata* and *Stramenopiles*. Representative organisms, lifestyle, and motility status are highlighted. Data points represent CPT values per organism.

We next investigated how these chanome size regimes group different types of organisms. The group with the least chanome investment (g1) includes parasitic protist species of the *Entamoeba*, *Leishmania*, *Cryptosporidium*, *Eimeria*, *Trypanosoma*, and *Plasmodium* genera; all lacking a nervous system (NNS). The parasitic myxosporean *T. kitauei*, a highly-derived class of *Cnidaria*, is also included in this group (**Fig. 2b,c**). A low CPT value in these organisms is consistent with the reduced complexification of ion channels observed in some protist parasites compared to free-living relatives^36^, and it is not simply explained by total genome reduction. Some species in this group have large genomes (e.g, *E. huxleyi*, 38,542 genes, 1.4 CPT; *T. cruzi*, 32,278 genes, 0.37 CPT) but encode fewer VGL-channels, as their relatives with smaller genomes do (e.g., *T. theileri*, 11,312 genes, 0.79 CPT). At the lower end of the second group (g2) we find the multicellular sponge *A. queenslandica* with a density of VGL-channels (1.63 CPT) only slightly higher than that observed in its unicellular g1 neighbors the coccolithophore *E. huxleyi* (1.4 CPT) and the diatom *F. cylindrus* (1.27 CPT). On the other hand, the unicellular choanoflagellates *M. brevicollis* and *S. rosetta*, and the multicellular placozoans, show CPTs between 2.7 and 3.5 and cluster next to the bulk of arthropod species, which are overrepresented in this same group despite their centralized brains (**Fig. 2b,c**). Group 3 (g3) largely contains vertebrate and invertebrate deuterostomes and includes a majority of organisms with centralized nervous systems (CNS). Notably, it also groups some organisms with a primitive, decentralized (e.g., *Nematostella*) or without a nervous system (e.g., *Paramecium*, *Stentor*) (**Fig. 2b,c**). Remarkably, the highest chanome investment group (g4) is characterized by early-diverging multicellular organisms with a diffuse neural net (NN) and unicellular protists lacking a nervous system altogether (cnidarians and ciliates) (**Fig. 2b-c**), indicating a disconnect between the availability of a vast electrically excitable toolkit and organismal complexity. This observation is reflected in the higher genomic chanome investment placed by cnidarians and ciliated organisms over bilaterians as a group (p<0.001, Wilcoxon signed-rank test), as well as over the median investment placed by vertebrates with complex brains (p<0.007, cnidarians; p<0.05 ciliated; Wilcoxon signed-rank test) (**Fig. 2e**). A focused analysis within protists further showed that extremes in total VGL-channel content may be associated with their capacity to move (**Fig. 2f**). The protists species with the most extreme CPT scores are also those that are motile (ciliates and flagellates) within both *Alveolata* and *Stramenopiles*, suggesting that the physiological imperative for motility may explain the extreme patterns of chanome investment within these unicellular eukaryotes.

### Global analysis of VGL-chanome composition variability

The diversification of channel types, and not just total VGL-chanome size, is relevant for the complexity and robustness of the signaling modalities available to a given organism^33^. Higher ion channel variability is considered to be indicative of greater computational and behavioral sophistication^31^. To analyze the distribution of channel types in each VGL-chanome, we classified channel candidates into (sub)families based on best-matching HMM hits in protein space. To study the patterns of chanome variability across organisms, we decomposed the total CPT score into subfamily-specific CPT values, a proxy for the genomic investment an organism places in the capacity to produce ion channels of different types (**Supplementary Fig. S3a**). We used these profiles to measure similarity in patterns of ion channel type investment among organisms and build a network connecting them accordingly (**Supplementary Fig. S3b,c**). By clustering this network we identified 12 clusters of organisms with similar profiles that we interpreted in terms of taxonomic groups, type of nervous system, and chanome investment (CPT groups) (**Supplementary Fig. S3c,d**). The network topology recovered clear associations between related organisms, as seen in a 2D network projection (**Fig. 3a**). The most evident topological segregation (vertical axis) distinguishes invertebrate (top) and vertebrate organisms (bottom), with 4 vertebrate clusters (green scale) primarily recovering mammals (c6), mammals and reptiles (c5), birds (c9), and fish (c2); and the remaining 8 clusters recovering nematodes (c12), arthropods (c11, c1, c3a), mollusks (c7), protists (c8, c4, c3b), and cnidarians (c10). Three clusters stand out as outliers based on their relationships with other groups of organisms. Most strikingly, the cnidarian cluster (c10), which also includes the only ctenophore examined (*M. leidyi*), localizes next to vertebrates in the bottom of the network, indicating greater chanome composition similarity with vertebrate than invertebrate organisms. Next, the nematode cluster (c12) localized apart from any other group of invertebrates, and with an orientation towards vertebrates, suggesting a qualitatively different chanome composition. Finally, one of the protist clusters (c8) localized apart from other protists and orienting towards vertebrates, suggesting changes in chanome composition in this subset of protists relative to others. Manually inspection of cluster membership associations of individual organisms identified additional surprising relationships. Sponges (*Porifera*) clustered within unicellular protists (c4), while unicellular choanoflagellates and placozoans clustered with mollusks in c7 and localized separate from unicellular organisms and closer to arthropods. These associations are explained by the presence of representatives of multiple channel types in choanoflagellates at low copy numbers (e.g., 1.78 copies per subfamily in average for *S. rosetta* versus 7.5 in human), despite their small VGL-chanome (32 channel candidates in *S. rosetta,* 2.9 CPT; vs, for example, 144 in mouse, 6.6 CPT). Notably, these unicellular organisms already include channels known to mediate the excitability of muscular and neuronal cells in animals (e.g., Na_v_ and Ca_v_) (**Supplementary Fig. S3e**). These observations further suggest a non-trivial relationship between electrically excitable toolkits and organismal and nervous system complexity.

**Figure 3.**
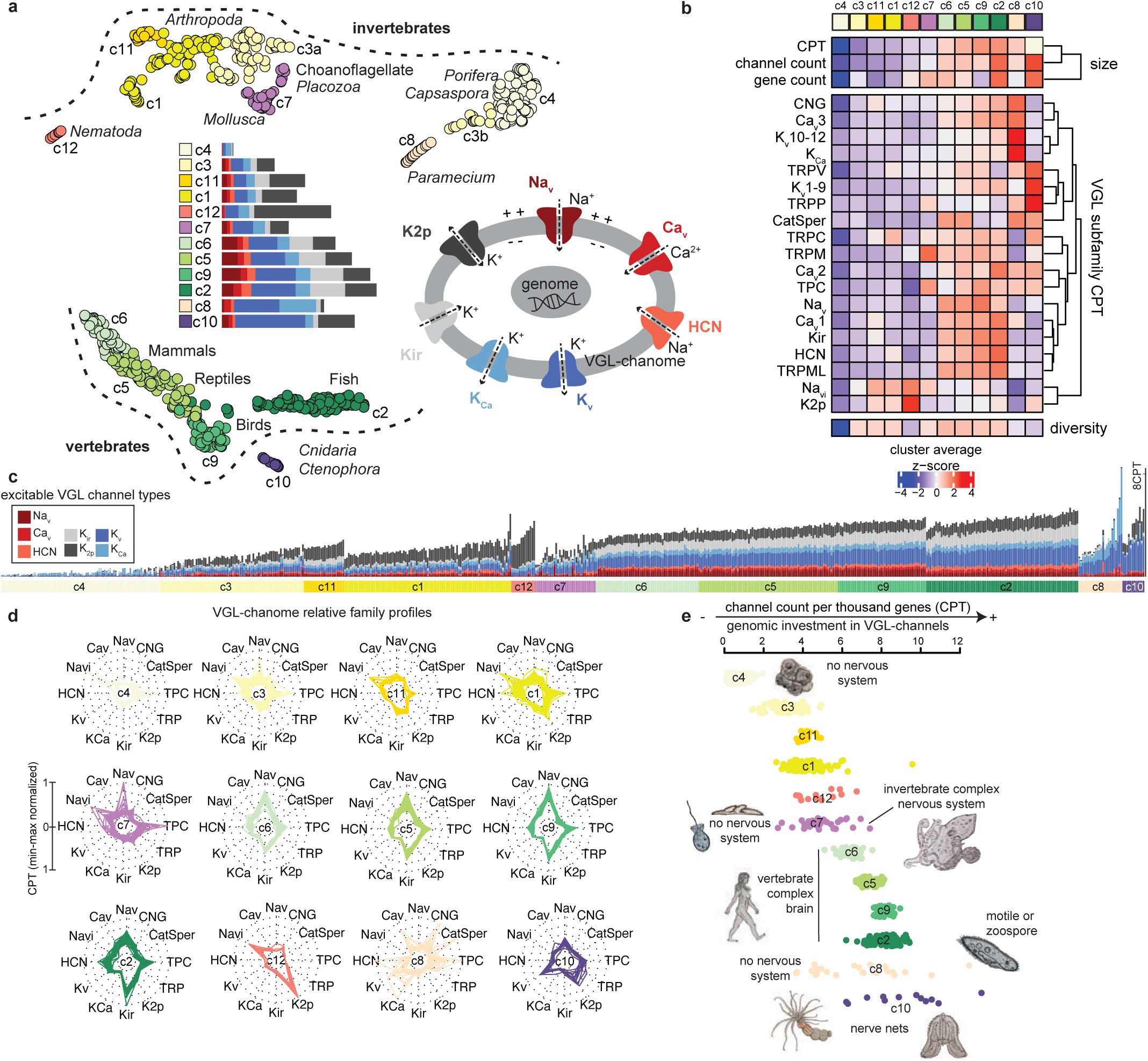
Global analysis of VGL-chanome compositional variability. **a)** 2-dimensional projection of organismal network connecting organisms (n=623) based on VGL-chanome composition similarity. Data points represent individual organisms. Colors represent clusters of organisms with similar VGL-chanome composition. Labels highlight cluster assignments and representative taxa and organisms. 2D coordinates were inferred using the Uniform Manifold Approximation and Projection (UMAP) algorithm. Clusters were defined using the graph-based walktrap algorithm. Bars insert shows VGL-chanome family composition profiles per cluster. Membrane insert illustrates the role of channel families in excitable membranes. **b)** Relative subfamily composition profiles. Data values represent row z-scaled size (upper) or CPT averages (bottom) per subfamily (row) and organismal cluster (column). **c)** VGL-chanome family composition profiles per organism. Data values represent CPT values per channel family and organisms considering only intrinsically excitable families. **d)** Multivariate radar chart depicting normalized VGL-chanome family composition profiles per cluster. Peaks visually highlight biases in family composition possibly related to lineage-expansions. **e)** Distribution of CPT values across organismal clusters. Data points represent individual organisms and are rank ordered within each cluster. Representative organisms, lifestyles, and nervous system traits are highlighted.

To better understand what differences in chanome composition underlie these associations, we estimated an VGL-chanome profile for each cluster and organism by aggregating family-based CPT scores (**Fig. 3a,c bars**); as well as per-cluster channel subfamily averages, genome and chanome sizes, CPT values, and relative family profiles (**Fig. 3b,d**). Clusters c8 and c10 have an unexpectedly large chanome (CPT and channel counts) relative to the size seen across invertebrate organisms (**Fig. 3b,e**). Protist cluster c8 in addition has a profile enriched in potassium (K_v_ 10-12, K_Ca_), Ca_v_, and CNG channels (c8); while the cnidarian cluster (c10) has one enriched in K_v_1-9 and a subset of TRP channels (**Fig. 3b**). In contrast, the chanome of the nematode cluster (c12) is more close in size to that seen in invertebrates (**Fig. 3b,e**), but very distinct in composition, with enrichment of K2p channels and depletion of Na_v_, TPC, and HCN channels (**Fig. 3a-d**). Na_v_, Kir, and HCN are in turn enriched in all vertebrate clusters (**Fig. 3b, Supplementary Fig. S3a,g**), which show cross-cluster chanome size variation with an increasing gradient from c6, c5, c9, to c2 (**Fig. 3b,e**; **Fig. 3a**, left to right). These patterns can be clearly appreciated at organismal level by inspecting the composition of representative species (**Supplementary Fig. S3e**).

To assess what types of ion channels are most determinant in clustering organisms, and thus likely varied systematically in subunit numbers within lineages during evolution, we trained a machine learning classifier to predict cluster labels based on subfamily profiles and estimate the relevance of each subfamily for the prediction task (Methods). The top 5 most informative subfamilies were K_v_1-9, K_v_10-12, Kir, K2p, and Na_v_ (**Supplementary Fig. S3f,g**). Thus selective expansions of three of the main families defining K^+^-dependent cellular excitability (K_v_, Kir, and K2P)^37^, together with expansions more broadly studied in Na_v_ channels^38^, seem to have been important determinants in the evolution and specialization of excitable cells. Notably, channels responsible for modulating membrane resting potential (K2P and Kir) and excitation mechanisms of action potential generation (e.g., K_v_, Na_v,_) are the primary determinants of chanome composition variation across protists and metazoans, and not those involved in sensory transduction and generator/receptor potentials (e.g., TRP and CNG) nor Ca_v_ channels. Estimated subfamily count and CPT profiles, network coordinates, and cluster annotations are reported in **Supplementary Table S1**.

### VGL-chanome diversity and the modes of expansion

Despite the wide range in VGL-chanome size across organisms, the genomic investment in VGL-channels alone does not seem to match phylogenetic relationships or perceived organismal or nervous system complexity. Instead, both chanome size and compositional variation determine (un)expected organismal relationships. Selective expansions of specific channel families are major contributors underlying these patterns, suggesting channel type preferences in the genomic investment of organisms. As chanomes expand or contract in size, they could do so equally across many or just a few of the channel families. A large and diverse chanome would imply having many genes encoding most of the channel types. A specialized one would be one with many genes of only a few channel types. To investigate the relationship between chanome diversification or specialization and size, we computed per organism a chanome diversity index measuring the deviation from a uniform distribution across subfamily copy numbers (Methods). Comparison of diversity and size revealed a remarkable pattern in which two main modes of expansion became apparent (**Fig. 4a**). VGL-chanome diversity follows a negative exponential relationship with size (CPT) in most organisms, increasing from small unicellular chanomes with low diversity to relatively small but diverse invertebrate chanomes and plateauing at large and diverse vertebrate chanomes (**Fig. 4a, Supplementary Fig. S4a**). Nematodes, cnidarians, and motile protists deviate from this pattern by having large chanomes of lower diversity. Mammalian-enriched clusters deviate in the opposite direction, thus having higher diversity than expected for their size (**Fig. 4b,c, Supplementary Fig. S4b**). Each of the three outlier groups of lower diversity presents a particular expansion in the number of channels of a given type. Organisms in the nematode or cnidarian clusters have, respectively, expansions of K2p or K_v_ channels, consistent with previous observations of K_v_ channel expansion in *Cnidaria*^39,40^. Organisms in the motile protists cluster have a general expansion of K^+^ permeable channels (**Fig. 4e**). Notably, organisms that departed from the growth pattern and yet parallelly acquired large but lowly diverse chanomes either have decentralized nervous systems or lack one altogether. Thus, not sheer chanome size but diversity seems to correlate with nervous systems complexity. Unlike those without or with diffuse nerve nets, organisms with a complex nervous system are characterized by a more diversified chanome investment (p<1e-05, Wilcoxon signed-rank test) but not necessarily a larger one (**Fig. 4d**). A pattern of specialized versus generalized diversity sets apart VGL- chanomes of organisms with complex nervous systems from those without (**Fig. 4e**).

**Figure 4.**
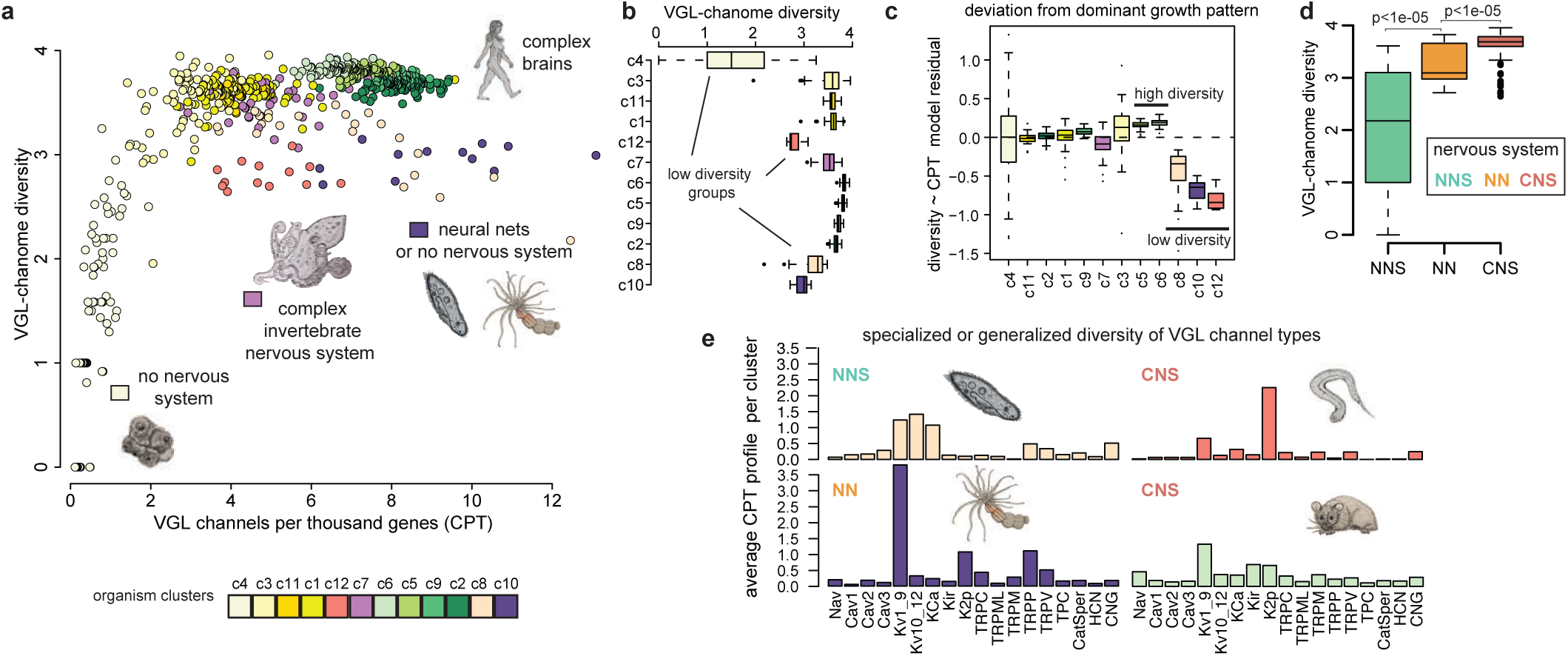
Patterns of VGL-chanome growth and diversity. **a)** Relationship between chanome size (CPT) (x-axis) and diversity (y-axis) highlighting organismal clusters (color-coded), representative organisms, and nervous system traits. Data points represent individual organisms (n=623). VGL-chanome diversity is measured using Shannon’s entropy. **b)** VGL-chanome diversity distribution across organismal groups. Data points represent individual organisms. **c)** Deviation from model predictions based on negative exponential relationship between VGL-chanome diversity and size (CPT) shown per organismal cluster (x-axis). Extremely high- and low-diversity clusters are highlighted. Data points represent individual organisms. Deviations are quantified based on model fit residuals (y-axis) (**Supplementary Fig. S4a**). **d)** Relationship between VGL-chanome diversity estimates and nervous system structural organization. Data points represent individual organisms. *P* values were calculated using Wilcoxon two-sided signed-rank tests. **e)** VGL-chanome subfamily compositional profiles of outlier organismal groups. Data values represent average CPT values per channel subfamily and organismal group.

### Cellular distribution of the VGL-chanome

Along with VGL-chanome growth and specialization, there are dramatic transformations in body size and cellular organization across metazoans. To investigate if and how such differences in organization impact the cellular distribution of the genomically available VGL-chanome, we re-analyzed single-cell transcriptomic atlases of a broad range of metazoans at whole-organism (Porifera, Placozoa, Ctenophora, Cnidaria, Nematoda) or multi-tissue (Arthropoda, Mammalia) level (**Fig. 5a**). We take transcription as an indicator of whether a cell type can use a given channel to regulate membrane voltage dynamics. We re-annotated major cell groups, including neuronal, glial, and muscle cells; as well as neuroid, neuroglandular, and peptidergic cells in early diverging metazoans, together with lineage-specific cell types (e.g., choanocytes in *Placozoa*, comb cells in *Ctenophora*, and cnidocytes in *Cnidaria*). We refer to cell types or groups generically as cell groups.

**Figure 5.**
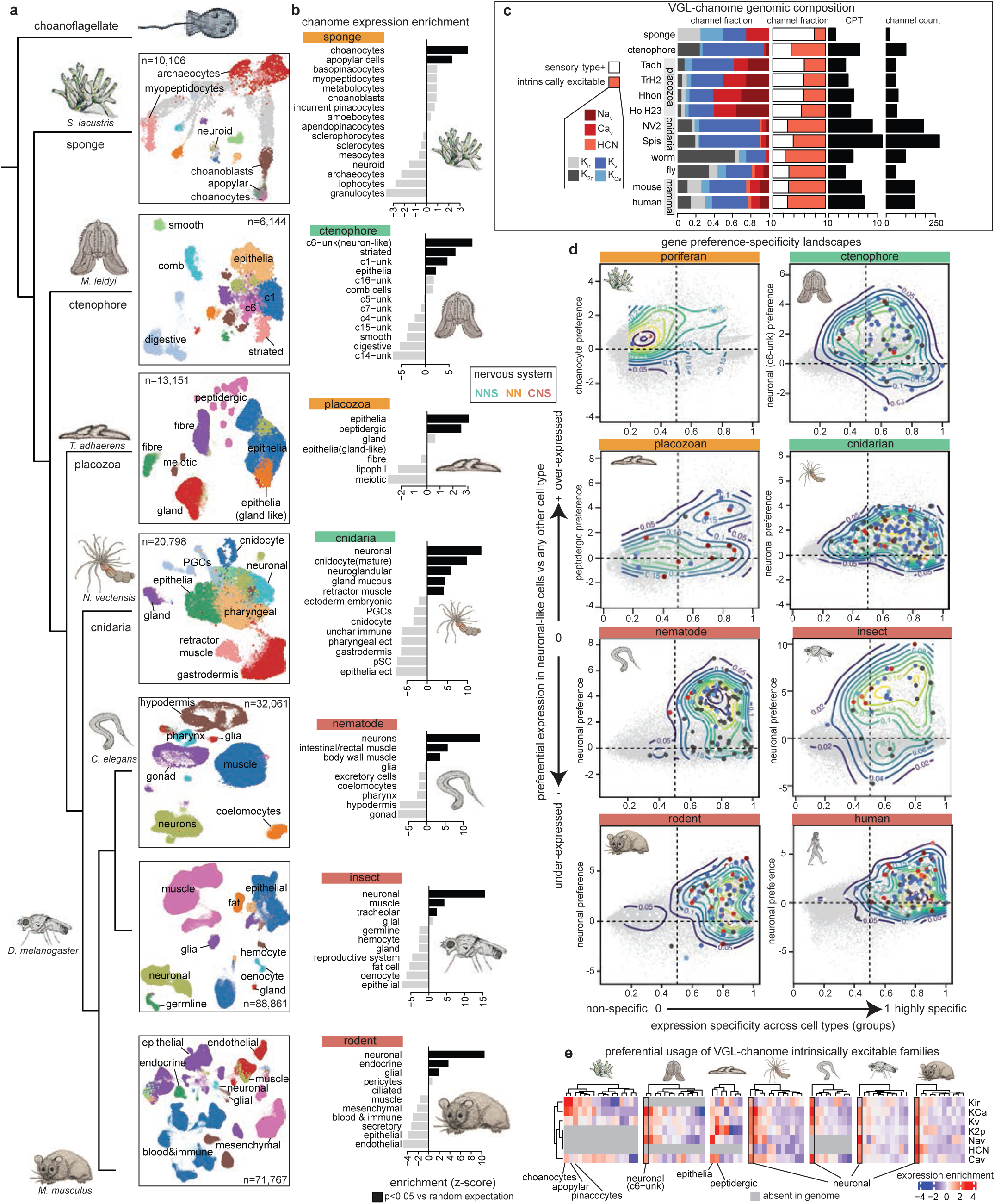
Cellular expression patterns of VGL-chanome. **a)** 2-dimensional representation of single-cell transcriptomic atlases. Data points represent individual cells labeled by major cell groups. 2D coordinates were inferred using the UMAP algorithm. Organisms are organised according to evolutionary relationships based on TimeTree estimates^99^. **b)** Preferential expression of VGL-chanome genes. Data values represent z-scores measuring the degree of deviation from random expectation of the average preferential expression values of VGL-chanome genes. Random expectation is estimated based on equally-sized random gene samples (n=10,000 samples with substitution). Unexpectedly high deviations are highlighted in black. *P* values were calculated based on gaussian null z-score distribution. **c)** VGL-genomic composition of representative organisms (rows). Channel family and type fractions are shown for intrinsically excitable- (left) or sensory-type- (TRP, TPC, CNG, CatSper, Navi) (middle) channels. VGL-chanome sizes are shown per organism in CPT and subunit counts (black bars, right). **d)** Preference-specificity landscapes representing the relationship between neuronal expression preference (y-axis) and gene expression specificity (x-axis). Data points represent individual genes. VGL-chanome genes are color-coded by channel family, and all genome background genes are shown in grey. Contour lines delineate areas of high density-based kernel density estimations over VGL-chanome genes. Choanocyte and peptidergic expression preference (y-axis) are shown respectively for Porifera and Placozoa, which lack neuronal cells. **e)** Average preferential expression values per VGL-chanome family (rows), cell group (columns), and organism (separate heatmap). Only intrinsically excitable families are considered. Data values represent pairwise preferential expression values computed separately across cell groups within each organism.

We found that VGL-chanome genes are preferentially expressed in subsets of cells in all organisms, including those without or with primitive forms of nervous systems, suggesting global restrictive expression of electrogenic ion channels across organisms irrespective of VGL-chanome size or composition, and independent of nervous system presence or organization (**Fig 5b,c**). This observation is supported by cell group and single-cell level analyses (**Fig 5b, Supplementary Fig. S5a**). We tested for cellular preferences at group level by calculating the average preferential expression values of VGL-chanome genes in each cell group and comparing this value with random expectation (Methods) (**Fig. 5b**). We identified groups that preferentially express the VGL-chanome genes (*P*<0.05, resampling) in all organisms. These groups are consistent with known excitable cells in human, rodent, insect, and nematode; and include neuronal, glial, endocrine, muscle, and germ cells^2^. Neuronal cells showed the strongest preference in all organisms with a nervous system. In ctenophore, this association highlighted a cell group originally reported as unknown (c6-unk)^41^. We found that this group also correlates with neuronal cells when comparing with transcriptomic signatures of *Nematostella, C. elegans,* and *Drosophila* (**Supplementary Fig. S5b**). We therefore interpret it as putatively neuronal (c6-unk neuron-like). In organisms lacking a nervous system, the VGL-chanome is preferentially expressed in placozoan peptidergic and epithelial cells, and in poriferan choanocytes and apopylar cells (**Fig 5b**). Single-cell level expression patterns further support these results. VGL-chanome expression levels and cell group annotations clearly overlapped in 2D cell landscape projections (**Supplementary Fig. S5a**). In most cases, expression patterns of the complete VGL-chanome or only the intrinsically excitable channel subset (Na_v_, Ca_v_, HCN, K_v_, KCa, Kir, K2p) overlapped, indicating consistency in the cell preferences of both sensory and intrinsically excitable channel types. This was not the case for the placozoan *Trichoplax*, where the latter were primarily expressed in peptidergic and not epithelial cells, suggesting segregated distribution of channel types among putatively excitable cells in this organism (**Supplementary Fig. S5a**).

A global analysis of gene expression preference and specificity patterns confirmed these observations and identified differences and similarities across organisms (**Fig. 5d-e, Supplementary Fig. S5**). We defined a gene preference-specificity space by comparing the expression signature of top excitable cells (choanocytes in *Porifera*, peptidergic in *Placozoa*, and neuronal elsewhere) versus gene expression specificity, and mapped VGL-chanome genes (family color-coded) to this space (**Fig. 5d**). The expression signature (y-axis) quantifies preferential expression in a specific cell group, while gene specificity (x-axis) quantifies the degree to which a gene is broadly (close to 0) or preferentially (close to 1) expressed across all groups (Methods). The more a gene localizes to the right side of this preference-specificity landscape, the more its expression is cell type specific. The more it localizes to the upper-part, the more it is neuron(-like)-specific. We observed a global neuronal expression preference in all organisms with a nervous system, as evidenced by an upper shift of the density of VGL-chanome genes (**Fig. 5d**). Notably, together with an upper shift, organisms with a complex nervous system also show a large rightward shift that is less pronounced in those with a more primitive decentralized nerve net (cnidarian and ctenophore), indicating that the VGL-chanome tends to be both generally specific and neuronal preferential in the former, while it may be more promiscuously used across cell groups in organisms with a primitive nervous system. In organisms without a nervous system, we observed either a slight preference but lack of specificity (poriferan) or partial preference and specificity (placozoan) (**Fig. 5d**). We confirmed a partial cellular specialization of the VGL-chanome in placozoans by re-analizing 4 whole-organism atlases (*T. adhaerens*, *Trichoplax sp, H. hongkongensis, and C. collaboinventa*)^42^ (**Supplementary Fig. S5c,d**). Both peptidergic and epithelial cells preferentially express the chanome in all cases. Most chanome genes are specific but have distinct preference patterns that result in multimodal landscapes (**Fig. S5d, Supplementary Fig. S5e**). In *Spongilla* (poriferan) we identified a few intrinsically excitable channels in the genome (n=4) (Kir, K_Ca_, K_v_, Ca_v_; one gene each). Most of its VGL-chanome is composed of subunits that match sensory type channels (e.g., 9 TRP, 2 CNG). Therefore, preferential expression in choanocytes is driven primarily by increased expression of only a few unique subunits (i.e., a four-fold increase of Kir and K_Ca_ genes) (**Supplementary Fig. S5f**). Although the second poriferan analyzed (*A. Queenslandica*) has similar chanome composition (**Supplementary Fig. S3e, Supplementary Table S1**), we did not find enough matching data in the available transcriptomic atlas ^41^ to verify VGL-chanome expression preferences.

In contrast to excitable groups of organisms without a nervous system, neuronal cells preferentially express a VGL-chanome composed of most channel families (**Fig. 5d, colors**). Family-wise differential expression analysis confirmed this pattern (**Fig. 5e, Supplementary Fig. S5g**). Neuronal cells in all organisms with a nervous system preferentially express most (if not all) types of intrinsically excitable channels relative to other cell groups, suggesting that increased expression of a diverse VGL-chanome may be a neuronal hallmark. Other non-neuronal, putatively excitable cells in these organisms show idiosyncratic specificity of some channel types (e.g., Na_v_ and K_Ca_ but not Ca_v_ in *Mnemiopsis* striated cells, Na_v_ and K_v_ and K_Ca_ but not HCN or Kir in *Nematostella* cnidocytes, and Kir in *Drosophila* glial cells (**Supplementary Fig. S5g**). Kir channels generally show increased expression in neuronal cells with an exception in *Drosophila* neurons, consistent with emerging reports of these channels as regulators of diverse physiological processes in several tissues of insects^43^. Putative excitable cells of organisms without a nervous system (sponge and placozoa) show increased expression of only some excitable channel types relative to other cell groups (e.g, K_Ca_ and Kir in *Spongilla* Choanocytes; K2p and K_v_ but not Na_v_ channels in *Trichoplax* peptidergic cells) (**Supplementary Fig. S5h**). Cell group absolute and preferential expression profiles and annotated single-cell atlases are reported in **Supplementary material**.

### Neuronal chanome specialization

Together with changes in body plan, metazoan evolution brought about remarkable transformations in nervous system structure. Neurons are the cellular units of all nervous systems, but some organisms organize them in a simple way, as in the diffuse neural nets of *Cnidaria* and *Ctenophora*, while others form complex subspecialized systems, as in worms or mammals with ganglia or brains. To investigate whether nervous system organization impacts neuronal VGL-chanome use, we re-analyzed single-cell transcriptomic data from the nervous systems of mouse^44^ and *C. elegans*^45^, and from all transcriptionally-identifiable neuronal cells of *Nematostella*^46^ (**Fig. 6**). We use these systems as models of high (rodent), medium (nematode), or low (cnidarian) structural and functional complexity.

**Figure 6.**
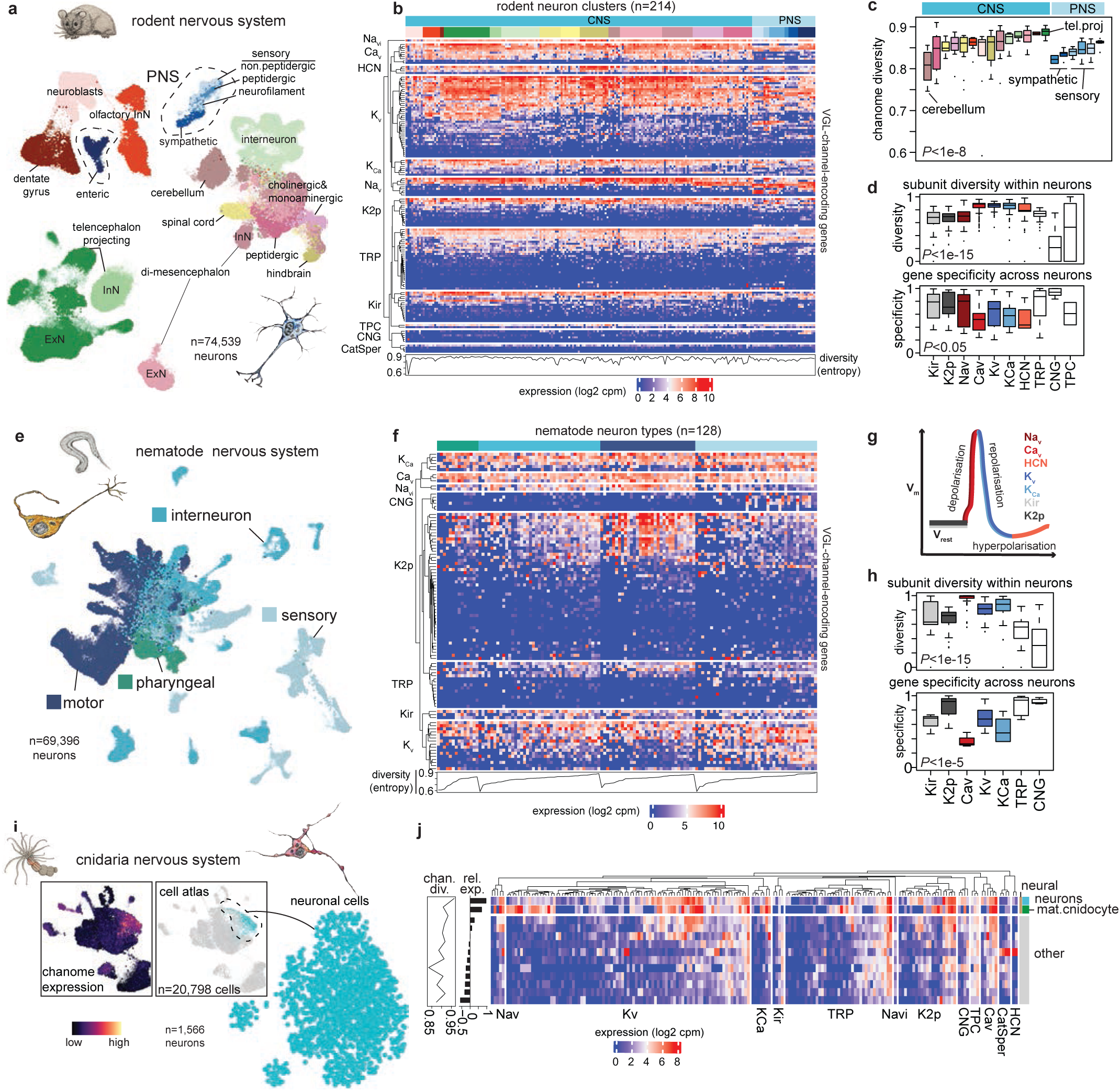
Neuronal VGL-chanome diversity across three different nervous system organizations. **a)** 2-dimensional representation of the single-cell transcriptomic atlas of neurons from the entire mouse nervous system. Data points represent individual cells labeled by neuronal taxonomic groups. **b)** Pseudobulk VGL-chanome expression profiles of 214 mouse neuron types (columns). **c)** VGL-chanome transcriptional diversity across taxonomic groups. Data points represent individual neuron types. Diversity is measured per neuron type using Shannon’s entropy over expression profiles. **d)** VGL-chanome family subunit diversity (upper plot). Data points represent individual neuron types. Diversity is measured per neuron type and channel family using Shannon’s entropy over subunit expression profiles. Subunit expression specificity (bottom plot). Data points represent individual subunit-encoding genes. Specificity is measured per gene and channel family using the Tau index. **e)** Same as (**a**) for neurons from the entire nervous system of *C. elegans*. **f**) Same as (**b**) for 128 neuron types of the nematode nervous system. **g)** Diagram illustrating the physiological role of the different VGL-chanome families in electrogenesis**. h)** Same as (**d**) across nematode neurons. **i)** 2-dimensional representation of neurons (blue) extracted from the whole-organism single-cell atlas of *Nematostella*. Inserts depict 2D representations of the whole atlas highlighting VGL-chanome expression (scale) and neuronal cells (blue). **j)** Pseudobulk VGL-chanome expression profiles of 13 *Nematostella* cell groups. Groups corresponding to neural cells (neuronal and mature cnidocytes) are labeled. Bottom black bars represent average relative expression values. Bottom line plot represents VGL-chanome transcriptional diversity values. 2D coordinates were inferred using the UMAP algorithm in all cases.

Rodent data include 214 transcriptionally-defined neuron types spanning 21 taxonomic neuron groups from 25 anatomical regions that sample both central (CNS) and peripheral nervous systems (PNS) (**Fig. 6a,b**). Comparison of pseudo-bulk profiles of taxonomic groups with profiles from non-neuronal cell types confirmed preferential expression of the VGL-chanome in all neuron groups (**Supplementary Fig. S6a**). Neurons express on average higher values of chanome-encoding genes than non-neuronal cell types (*P*<1e-06, Wilcoxon signed-rank test), with higher levels in both CNS and PNS relative to other cells (*P*<1e-04, Kruskal–Wallis test) (**Supplementary Fig. S6b**). To contrast overall chanome diversity across neuron types and classes, we estimated a transcriptional diversity index per neuron type (Methods). A higher diversity indicates a more uniform expression of VGL-chanome genes, irrespective of channel type. We found lower diversity in PNS versus CNS neurons (**Fig. 6c, Supplementary Fig. S6c**), and differences in diversity across taxonomic groups (*P*<1e-08, Kruskal–Wallis test) (**Fig. 6c**). Within CNS neurons, cerebellar neurons (light brown) showed the lowest diversity and telencephalic projecting excitatory neurons (*tel.proj*, dark green) the highest. Despite their overall lower diversity, PNS neurons also displayed a diversity gradient, with enteric neurons having the highest diversity, followed by sensory neurons, and with sympathetic neurons having the lowest. Thus, in a highly-complex and specialized nervous system, neuronal VGL-chanome diversity varies across structures, possibly reflecting functional subspecialization and differences in neurocomputational potential.

VGL-chanome diversity is achieved at 2 levels: by the combinatorial expression of different channel types, and of multiple subunits per type. In rodent, some channel types seem to express many subunits across most neurons (e.g., Ca_v_), while others just a few, and with different degrees of specificity (e.g., Na_v_) (**Fig. 6b**). To quantify these patterns, we measured diversity at subunit level within neuron types, as well as subunit specificity across neurons and found cross-family differences (*P*<1e-15 diversity, *P*<0.05 specificity; Kruskal–Wallis test) (**Fig. 6d**). Channels impacting resting membrane potential (Kir, K2p) showed lower diversity and higher specificity than those involved in de-, re-, or hyperpolarization (**Fig. 6g**), suggesting use of few of the available subunits with differential patterns in some neuron types. Among depolarizing channels, Na_v_ are the exception, having relatively low diversity (median=0.7) and high (median=0.8) but variable (IQR=0.5) specificity, suggesting preferential expression of a subset of subunit genes in specific neuron types (e.g., Na_v_1.7, Scn9a; and Na_v_1.9, Scn11a in sensory neurons) together with broad expression of others (e.g., Na_v_1.6, Scn8a; and Na_v_1.3, Scn3a in most neurons). In contrast, Ca_v_-encoding genes showed the highest diversity (median=0.89) and lowest (median=0.52) and least variable interneuronal specificity (IQR=0.24), indicating that most available subunits are broadly expressed across neurons. Depolarizing families (K_v_ and K_Ca_) showed high diversity (median=0.87 in both) and intermediate specificity (median=0.68 in K_v_ and 0.57 in K_Ca_), indicating that most neuron types use relatively large and similar subunit complements of these channels. Among sensory type channels, TRP showed intermediate diversity (median=0.74) and high (median=0.87) but variable (IQR=0.39) specificity. CNG channels are low-diverse (median=0.21) and particularly specific (median=0.94), with preferential expression in PNS.

Nematodes have lower structural diversity in both the nervous system and the VGL-chanome (e.g., absence of Na_v_ and HCN) than rodents. We found that these differences impact some aspects of neuronal VGL-chanome use but not others. Transcriptomic data of 128 worm neuron types, including sensory, motor, pharyngeal, and interneuronal cells, confirmed overall preferential chanome expression in neurons relative to non-neuronal cells (**Fig. 6e,f; Supplementary Fig. S6d**) and variable diversity across neuron classes (*P*<0.05 Kruskal–Wallis test) (**Supplementary Fig. S6e**). The VGL-chanome of sensory neurons was the most diverse (median=0.82), followed closely by motor and interneurons (median=0.81). Pharyngeal neurons showed the least diversity (median=0.77) (**Supplementary Fig. S6e**). Diversity ranks within functional classes uncovered several notable observations (**Supplementary Fig. S6f**). The most diverse chanomes within sensory and motor neurons correspond to the AWA and RMD neuron types, both known to have a polymodal function -- i.e., encode several sensory stimuli and/or display both sensory and interneuron function. These two neuron types are also among the few known to fire all-or-non regenerative action potentials^47,48^. Certain subunit diversity and specificity patterns set apart nematode and rodent neuronal chanomes. In nematode neurons we observed differences between families of K^+^ channels involved in either resting potential (K2p and Kir) or repolarization (K_V_, K_Ca_) (**Fig. 6h**). K2p channels are low-diverse (median=0.72) and highly specific (median=0.92), with a visible enrichment in motor neurons (**Fig. 6f**). This pattern may be associated with subfunctionalization of this lineage-expanded ion channel family. K_Ca_ are relatively more diverse and less specific than K_v_, while Ca_v_ are similarly highly diverse and specific as in rodents (**Fig. 6b**). High and diverse Ca_v_ expression goes beyond neurons, including additional non neuronal excitable cells (**Supplementary Fig. S6d**). The broad use of Ca_v_ and K_Ca_ subunits possibly reflects the dominant electrogenic role for Ca^+2^ ions in this organism which lacks Na_v_ channels.

Unlike nervous systems with complex structural organization, in the simpler diffuse nerve net of cnidaria we were not able to confidently identify and annotate distinct transcriptional neuron types. The *Nematostella* single-cell atlas includes 1,566 annotated neurons that preferentially express VGL-chanome genes (**Fig. 6i**). Independent analysis of these neurons revealed a relatively homogenous neuronal landscape, with low modularity relative to the structured 2D neuronal landscapes of rodent or nematode. To verify this qualitative observation of neuronal transcriptional homogeneity in *Nematostella*, we measured pairwise transcriptional distances within each organism following a subsampling strategy (Methods) (**Supplementary Fig. S6g**). This strategy estimated significantly larger interneuronal distances in nematode and rodent relative to those seen in cnidarian neurons, thus confirming our observation. We therefore limited our neuronal chanome analysis to only one major neuronal group in this organism. Comparison of neuronal chanome expression with non-neuronal cell types confirmed higher relative expression in neurons and evidenced a close relationship with mature cnidocytes, which also express high levels of chanome genes (**Fig. 6j**). *Nematostella* neurons and cnidocytes share progenitor cells and a terminal selector gene (POU4) that has been shown to be required for ion channel expression^49^. In a second cnidarian (Hydra), both cell types also share progenitors and are considered sistercells^50^. The low neuronal heterogeneity, high diversity, and the large number of ion channels in both neural cells (neurons and cnidocytes) is consistent with their reported lack of definite cell fate specification and high plasticity in *Nematostella*^50^.

### Relationship between chanome composition and electrophysiological properties in a simple cnidarian

How are physiological functionality and VGL-chanome composition linked? To investigate how these two aspects play together in early-diverging metazoans, in which neuronal excitability and coding are not well-understood, we made use of the cnidarian model organism *Nematostella vectensis* (**Fig. 7a**). Nematostella has three classes of neural cells: sensory and ganglionic neurons and cnidocytes^51^. We could clearly distinguish transcriptionally only one group of neurons, however (**Fig. 6j**). Therefore, to explore how the expression of ion channels in a subset of identified neurons relates to its excitability, we used a neuron-specific transgenic line (*NvElav1::mOrange*)^52^ (**Fig. 7b,b’**). We isolated mOrange positive neurons from the tentacle and performed whole-cell patch clamp electrophysiology to describe basic excitability properties (**Fig. 7c,d**). Nematostella has highly specialized neurons with elaborate morphology and long protruding neurites (**Fig. 7b’**), suggesting the need for a rapid and long-distance signaling mechanism. In agreement with this view, and consistent with previous observations^53,54^, we found that the isolated neurons can produce fast spikes, reminiscent of mammalian sodium-mediated action potentials (APs) (**Fig. 7d**). To understand the composition of these all-or-none electrical events, and subsequently relate their biophysical features with VGL-chanome expression, we performed patch-clamp experiments describing the current composition and voltage responses to depolarizing current injections in elav-positive cells (**Fig. 7e-g and Supplementary Fig. S7a,b**).

**Figure 7.**
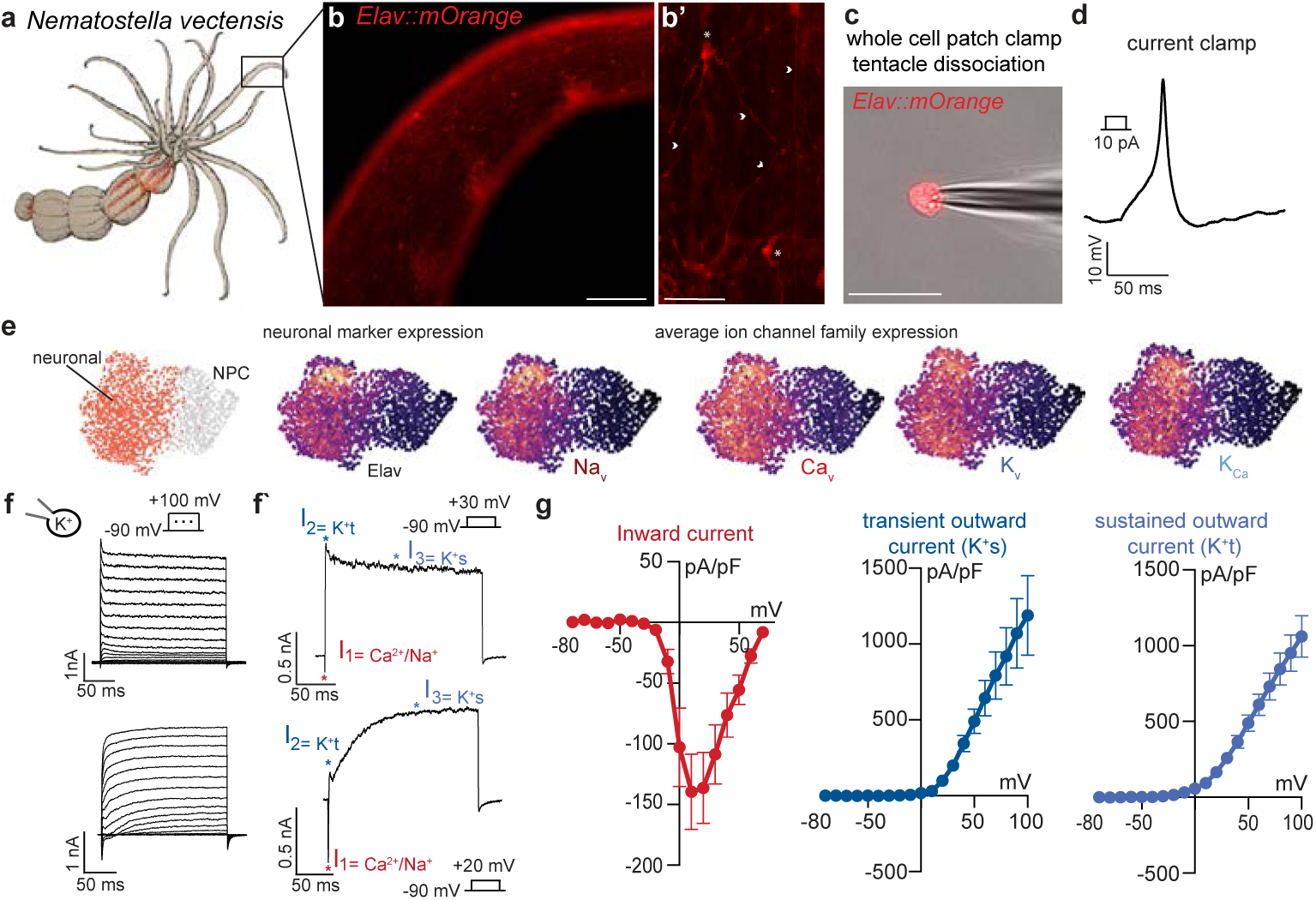
Nematostella has a nerve net containing neurons that produce overshooting sikes. **a)** The burrowing sea anemone *Nematostella vectensis* has a diffuse nerve net **(b)** here shown by labeling through a transgenic line (*Elav:1:mOrange*) in the tentacles (scale bar 200 um) with **(b’)** complex neuronal shapes and long extending neurites (annotation: star= cell body, arrow= neurites, scale bar 50 um). **(c)** Acute dissociations can be made from the tentacles and used for whole cell patch clamp recordings to describe electrical properties (scale bar 50 um). **(d)** Example of overshooting spike after injection of a 10 pA current. **(e)** *Elav1* positive neurons can be clearly identified in single cell data and contain VGL-chanome components similar to mammalian neurons. **(f)** Example traces for current voltage relationships and **(f’)** individual traces showing the different current components. **(g)** Quantification of current/voltage (IV) relationship for inward current (n=8), transient outward current (n=6) and sustained outward current (n=9). Currents corrected for cell size (pA/pF).

Whole-cell currents were recorded using a voltage step protocol (**Fig. 7f and Supplementary Fig. S7a,b**). The current composition was quantified as described in f’ considering inward current and sustained and transient outward currents (**Fig. 7g**). We observed different generic types of whole-cell currents in our recordings (n=9) (see, for example **Fig. 7f and f’)**, reflecting distinct electrophysiological cell types or states (also reflected in distinct spiking patterns - **Supplementary Fig. S7a and b**). In general, inward currents activated around -20 mV with peak amplitude around +10 mV, similar to currents reported in early studies of *Cyanea capitella* neurons^54^, confirming that cnidarian neurons have a positive voltage shift with unknown reason. This activation range is, however, close to the AP threshold measured to be at -11 mV (+/- 0.564 mV) (**Supplementary Fig. S7c**). Outward currents were diverse and complex, but two distinct currents could be quantified in most cells: a transient outward current (K+t) at the beginning of the voltage step (most likely an A-type current), and a sustained current (K+s) (**Fig. 7g**).

Nematostella has large genomic channel expansions (**Fig. 4d**)^39,40^. In our analysis we find that the majority of these expanded excitable channels are expressed in neural cells (neurons and mature cnidocytes) (**Fig 6j**). To understand how the cellular usage of the VGL-chanome relates to physiological function in a specific neuronal subpopulation, we used an *in silico* cell-sorting strategy to enrich for *elav1*-expressing cells based on transcriptome evidence from *Nematostella’s* single-cell atlas (**Fig. 7e** and **Supplementary Fig. S7g).** Among cells annotated as neuronal, we isolated those with the strongest evidence of *elav1* expression by sub-clustering and differential expression analysis. We detected expression of 98 out of 134 VGL-chanome genes encoding subunits of intrinsically excitable channels in these cells, including 13 out of 20 described K_v_ NveShaker channels (**Supplementary Fig. S7g,h**). Some Nematostella shaker channels have been reported to have unusually high voltage activation thresholds in heterologous expression systems, producing visible currents only around -10 mV^39^. This activation threshold matches the inward current peak, the outward current activation range, and the AP threshold of -11 mV in our measurements (**Supplementary Fig. S7c**). These channels are therefore well-suited to mediate AP repolarization in Nematostella neurons, similar to bilaterian shaker channels^55^. With respect to depolarizing channels, *elav1* cells express 4 out of the 5 described Nematostella Na_v_ channels (three nonselective) (**Supplementary Fig. S7h**)^56^, and 6 Ca_v_ channels. Therefore, both expression data and electrophysiological recordings suggest that *elav1*-positive neurons of the tentacle are both functionally and transcriptionally diverse, with complex inward and outward currents likely produced by a diverse combination of VGL-channels.

Complementary current-clamp experiments demonstrated that Nematostella neurons can produce two generic types of excitation events (graded potential and all-or-nothing action potentials) (**Supplementary Fig. S7**; cell 1, **S7a’** and 2, **S7b’**), consistent with observations in the motor neurons of *Aglantha digitale*^57^. With an average width of 7.971 ms (+/- 1,387 ms, SEM) (**Supplementary Fig. S7c)**, these spikes were significantly slower than APs recorded in some mammalian neurons (∼0.22 ms)^58,59^. Irrespective of the type of voltage change, *elav1* cells produced only one single spike, pointing towards a strong depolarization block. The two above-mentioned characteristics (spike width and depolarization block) suggest that the inward current could be mainly carried by calcium, mediated either by the nonselective Na_v_ family described in cnidarians^56^ or by canonical Ca_v_ channels.

## Discussion

The origin and evolutionary history of nervous systems has puzzled biologists for centuries^4,9,60–62^. Modern efforts investigate the evolution of neuronal cells by tracing developmental lineages or “neuronal genes” across model organisms and early-diverging metazoans^42,63–69^. Here we take a different perspective and focus on the molecular system that endows neurons with their unique electrical properties: the VGL-chanome. The electrical impulse is the elementary unit of nervous system function^8^. But electrical signaling is a preneuronal feature employed by a diverse set of excitable cells that do not share a developmental origin or evolutionary history^2^. They all share instead the ability of using ion channels to actively modify membrane potential for signaling. Neurons extend this ability by combining analog and digital signals to perform complex computations as part of information processing networks. By characterizing the voltage-gated ion channel complements (VGL- chanomes) of 623 organisms with different degrees of behavioral and structural complexity, dissecting their expression patterns in 11 whole-body cell atlases and 3 entire nervous systems, and recording electrical properties of “primitive” neurons in the sea anemone, we uncovered patterns of VGL-chanome evolution and function that may have enabled the emergence of neuronal excitability and nervous system complexity.

### Parallel VGL-chanome evolution

Our data highlight a disconnect between the genomic availability of ion channels and organismal or nervous system complexity. Unicellular organisms with complex behaviors (e.g., ciliates), early-diverging metazoans with nerve nets (e.g., cnidarians), and vertebrates with complex brains (e.g., mammals) have all evolved vast VGL- chanomes. This result matches early paradoxical reports of protozoa having more channels than humans (e.g., K^+^ channels in Paramecium)^70^. By analyzing 124 protist genomes, we revealed an association between a vast chanome and an active lifestyle. A large VGL-chanome in free-living motile protists agrees with the remarkable electrophysiological and behavioral complexity^36,71–74^ that has earned them the recognition of “swimming neurons” and tractable models for integrative neuroscience^20^. These humble organisms have evolved active membrane properties reminiscent of neurons, allowing them to combine sensory and motor functions in a single cell. Early-diverging metazoans with diffusive nervous systems (ctenophores and cnidarians) also evolved expanded VGL-chanomes similar in size and composition to vertebrate bilaterian chanomes. These results are consistent with previous reports of convergent ion channel content in the major lineages of animals with nervous systems^29^. We suggest that neuron-like unicellular organisms, organisms with diffuse nerve nets, and organisms with highly-complex centralized nervous systems, all acquired vastly expanded VGL-chanomes independently via gene-family expansions. A vast VGL-chanome is therefore not a signature of organismal or nervous system complexity.

### Two modes of VGL-chanome evolution

Our genomic analysis uncovered two modes of evolution in which VGL-chanomes acquired either a diversified or a specialized channel-family profile. Ion channel family composition is important for the complexity of electrical signals available for a cell^32^. Within protists and non-bilaterians VGL-chanomes specialized. Motile protists disproportionately acquired K^+^ channels, while non-bilaterians with diffuse nerve nets expanded K_v_ channels. In contrast, bilaterians with highly-complex nervous systems (e.g., mammals) acquired diversified VGL-chanomes with a more uniform representation of most channel families. Expansions towards lower diversity occurred only in organisms that either have a decentralized nervous system or lack one altogether, and are consistent with reported gene family expansions^39,40,70^. Nematodes are a bilaterian exception that expanded K2p channels while losing other families^75^. Peculiar physiological adaptations in these animals, like small cells^76^, high membrane resistance^77^, or preferential use of analog over digital signaling ^48^; might explain the smooth operation of their nervous system despite a contracted and specialized VGL-chanome. The acquisition of a diversified chanome in vertebrates was likely facilitated by a major genomic loss event in the ancestor of deuterostomes, followed by bouts of gene expansions^29^. This genomic contraction also correlates with intermediary nervous system simplification in stem chordates like tunicates. A second reported loss event in the ancestor of Arthopoda and Nematoda (ecdysozoans)^29^ might explain the compact chanome of the former and the subsequent lineage changes in the latter. Taken together, our data support a scenario of independent evolution of neuron-like unicellular organisms and diverse nervous systems that use similar ancient molecular tools to mediate excitability. Repeated bouts of elaboration and simplification of nervous systems associated with expansions and contractions of the VGL-chanome likely played a role. The evolution of highly-complex nervous systems was not a stepwise progression of expanding complexity, and it involved the acquisition of a diverse VGL-chanome.

### Neurons recruit a diverse VGL-chanome

The onset of multicellularity allowed for cell type specialization and growth but also created the need for long-range coordination. Neuronal systems specialized for electrical signaling are the solution to this problem. How are the cells composing these systems distinct from other excitable cells? We found that all organisms distribute their VGL-chanomes selectively across a few cell types that coincide with known excitable cells in nematodes, insects, rodents, and mammals; supporting the view of VGL-channel expression as diagnostic of excitability^2^. We detected selective expression in nerveless organisms too. The preferential expression of the VGL-chanome of sponges in choanocytes and apopylar cells, and expression of certain channels in pinacocytes (e.g., K_v_), suggests it mediates ciliary beating, contractility, and sensory perception in these organisms^67^. Placozoan VGL- chanomes are preferentially used in peptidergic and epithelial cells, but with distinct preferences (Na_v_-like in epithelial, K^+^ in peptidergic, and Ca_v_ shared). Importantly, we do not find cell types in nerveless animals overexpressing all channel families known to mediate intrinsic excitability. These observations are consistent with a scenario in which extant nervous systems derived independently from excitable proto-nervous tissues that use VGL-channels to directly mediate simple behaviors in stem animals^29,78^. In striking contrast, neuronal cells in all organisms overexpress a diverse set of excitable channel types, suggesting that their unique electrical specialization and neurocomputational capacity is not the result of the presence, amplification, or biophysical tuning of a particular channel family, but rather the combinatorial expression of subunits of most (if not all) intrinsically excitable channel families. Non-neuronal excitable cells show idiosyncratic channel type preferences, consistent with the view that different categories of excitable cells use their own regulatory elements to select genes that endow them with specific kinds of electrical excitability^2^. Such restrictive expression could also have a role in allowing structural changes in individual channel proteins due to lower pleiotropic constraints, thus facilitating tissue-specific physiological adaptations^31,79^. In the context of multicellular division of labor, where the formation and maintenance of plastic neural circuits enables information exchange between cells^69^; information coding, fast transmission, and network resilience may benefit from the simultaneous availability of many different types of ion channels. Coexpression of functionally overlapping ion channels allows neurons to be simultaneously resilient and plastic, withstanding perturbations and rewiring while maintaining functionality^33^.

### VGL-chanome tuning and nervous system specialization

Our single-cell analysis indicates that nervous system complexity is reflected in the degree of neuronal transcriptional heterogeneity. Yet, all neurons express diverse VGL-chanomes. Cell intrinsic and network properties within nervous systems impose biophysical constraints to ion channel function and information coding capacity. Our data suggests that the physiological implications of these constraints are reflected in VGL- chanome composition and usage. Neurons embedded in networks (e.g., CNS and enteric in rodents, and neurons in cnidarian) express more diverse VGL-chanomes than specialized sensory cells of the PNS, consistent with the imperative for degeneracy and resilience in the former^33^. Expression of a specialized VGL-chanome may allow for adaptations (e.g., unique sensory modalities or toxin resistance), but also implies higher vulnerability upon perturbations. The lower resilience of a VGL-chanome of low degeneracy or functional overlap is likely more consequential in a network than a sensor. A genomically diverse VGL-chanome provides the tools to implement both strategies via regulation, thus fitting the functional needs of a diversified nervous system. Our data on *C. elegans* neurons also suggest an association between VGL-chanome usage and function. Neurons that are multi- or polymodal (e.g., AWA and RMD) and are able to fire all-or-none action potentials, show the highest diversity. Information processing via both graded sensory input and digital signals in these neurons implies enhanced neurocomputational properties that may be associated with VGL-chanome diversity.

Expanded and specialized VGL-chanomes may support neuronal subfunctionalization. Some channel families contribute to neuronal transcriptional flexibility more than others, and their relative contributions seem to correlate with the genomic changes observed during evolution. Our data highlight a role for K^+^ (K_v_, Kir, and K2P) and Na_v_ channel expansions in excitable cell evolution. K2p channels are expanded in nematodes, and their subunits are also more specifically used across neurons in *C. elegans*. Na_v_ are expanded in vertebrates, and their subunit usage combines broad and highly specific expression in rodent neurons. The majority of expanded K_v_ in cnidaria are expressed in one of the two neural cell types. Genomic variability of Ca_v_ channels across metazoans is much lower, and their subunits are expressed jointly and very broadly in all organisms, suggesting strong pleiotropic constraints. Evolutionary constraints on Ca^+2^ channel usage in electrical signaling may be associated with the cytotoxicity of this ion and its ancient and ubiquitous role in several intracellular signaling^2,80,81^. The reasons why specific types of channels expanded and not others in different lineages remain unknown. Neuronal subfunctionalization possibly played a role, along with historical contingency, genomic, biophysical, and physiological constraints.

Our general understanding of the constraints under which neurons and ion channels function, and its relation to genome evolution, is very limited. Conclusions are commonly drawn from a few model organisms. Invertebrates often have to cope with additional biophysical challenges, like the use of very small neurons and/or calcium as main inward current carrier^47,48,82,83^. Investigating the biophysical implications of genome evolution in a broad range of animals will help clarify the parallel origins of nervous system functions. The characterization of expanded K_v_ and Na_v_ channels of stem animals and their similarities with those in vertebrates are notable efforts in this direction^39,56,84^. Nematostella has an expanded and specialized VGL-chanome that is preferentially expressed in neural cells (e.g., neurons, cnidocytes). Neurons are largely transcriptionally homogeneous and express many subunits of most families. Cnidocytes are considered cellular innovations that characterize cnidarians. Regulated expression of expanded K_v_ subunits might have contributed to the individuation of these neuronal sistercells. The low transcriptional heterogeneity and large number of ion channels in both neural cells is consistent with their reported lack of definite cell fate specification and high plasticity in Nematostella^50^. Our electrophysiological recordings of cnidarian *N. vectensis* confirmed that their independently evolved neurons generate fast inactivating inward currents and spikes similar to mammalian sodium-mediated action potentials (APs). However, these are slower, single spikes, presumably driven by calcium or mixed inward currents possibly mediated by nonselective Na_v_ channels^56^. Morphological and molecular studies suggested heterogeneity of neural cell types^51^. We observed heterogeneity in electrophysiological behavior despite apparent transcriptional homogeneity, with complex inward and outward currents likely produced by a diverse combination of VGL- channels, suggesting interindividual neuronal variability and channel usage promiscuity.

Finally, considering patterns of genomic expansion, collective overexpression, and subunit diversity, we propose that the evolution and cell restricted recruitment of a diverse VGL-chanome constitutes a neuronal hallmark. Regulatory tuning of such a chanome across specialized neurons may contribute in shaping the signals and responses of complex nervous systems and maintaining their robustness and plasticity. It is an exciting time to revisit old questions in light of new molecular data and technical advances that now allow in-depth empirical analysis and large-scale comparisons. Our integrative study provides new insights into the interplay between excitable molecules, excitable cells, and nervous system evolution and reports a valuable resource for future structure-function investigation of ion channels and excitable cells.

### Limitations of the study

The lack of data availability for ctenophores and poriferans precluded reproducible analyses of VGL-chanome composition or preferential expression. These organisms should be explored in more depth once new genomes, gene annotations, and single-cell atlases become available. Our genomic analysis primarily focused on mapping patterns of global VGL-chanome change across many taxa. Phylogenetic analysis of specific taxa from our curated resource may further clarify the evolutionary dynamics of lineage-specific channel expansions and contractions.

## Supplementary Figures

**Supplementary Figure 1.**
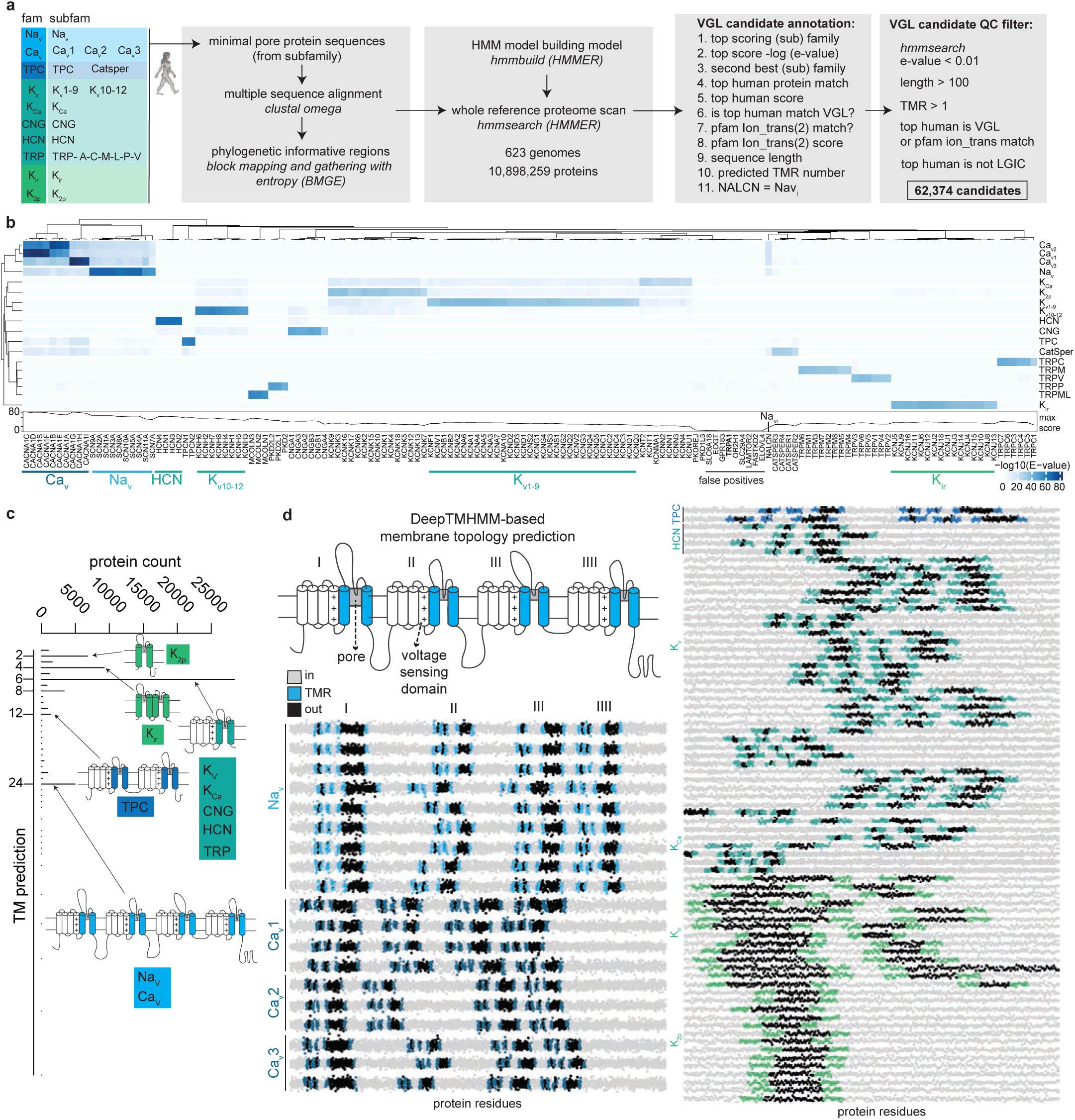
VGL-chanome identification and annotation workflow. **a)** Workflow to identify, annotate, and curate VGL-chanomes from genome (proteome) reference sequences. **b)** Probabilistic matching scores for human proteins (columns) identified as potential ion channels based on searches using HMM ion channel subfamily models as query (rows). **c)** Distribution of the number of predicted transmembrane segments (TMR) across 62,374 VGL-channel candidates from 623 genomes. Predictions are based on the DeepTMHMM model. **d)** Visualization of membrane topological predictions of a subset of human VGL-channels. Each data point represents protein residue (amino acid). Segments are color-coded according to predictions: In, intracellular; out, extracellular; TMR, transmembrane region. Top panel illustrates the known topology of 24 TMR domain ion channels shared by Na_v_ and Ca_v_ channels.

**Supplementary Figure 2.**
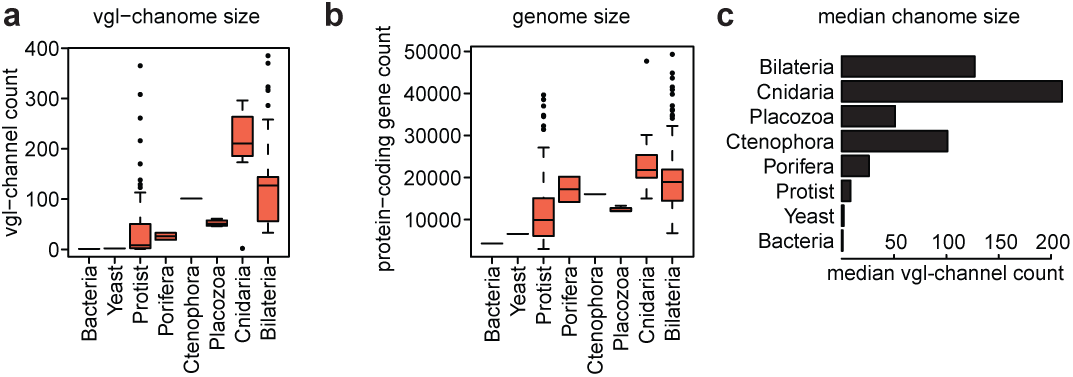
VGL-chanome and genome size statistics. **a)** VGL-channel count distribution across taxa. Data points represent individual organisms. **b)** Protein-coding gene count distribution across taxa. Data points represent individual organisms. **c)** Median VGL-channel counts across taxa.

**Supplementary Figure 3.**
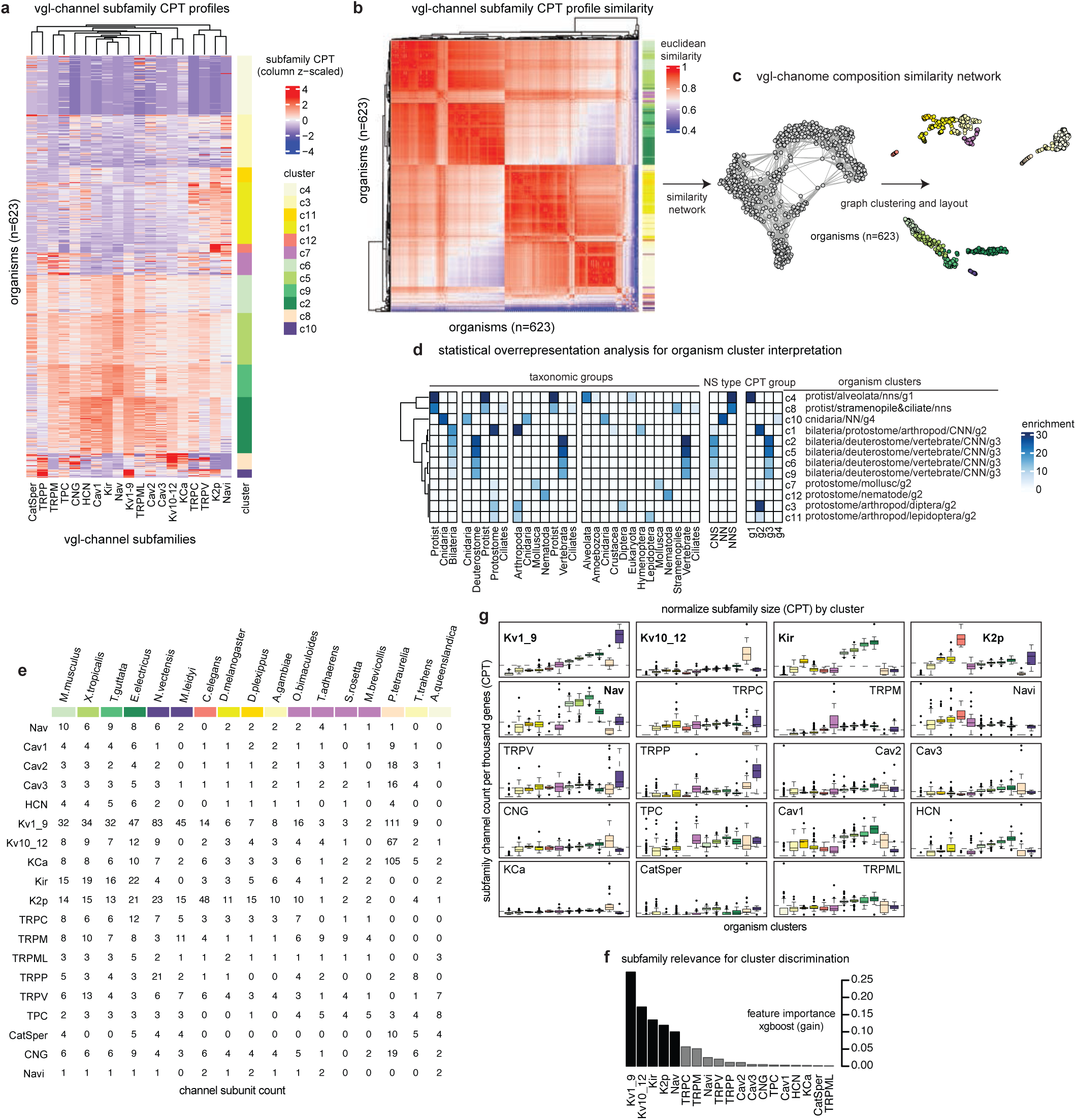
VGL-chanome compositional variability. **a)** VGL-channel subfamily (columns) profiles per organism (rows) organized by organismal cluster. Data values represent columnwise z-scaled subfamily CPT values. **b)** VGL-channel subfamily profile similarity. Data values represent Euclidean distance between pairs of subfamily profiles. **c)** Network connecting organisms based on VGL-channel subfamily profile similarity and illustrating clustering approach. **d)** Statistical overrepresentation analysis of taxonomic groups, nervous system organization traits, and CPT investment groups (columns) within organismal clusters (rows). Data values represent -log10(*P*) values. *P* values were calculated using binomial tests. **e)** Subunit counts per best-match subfamily (rows) for representative organisms (columns). **f)** Subfamily relevance for organismal cluster discrimination. Data values represent feature importance (gain) scores computed using xgboost. **g)** Distribution of VGL-chanome subfamily sizes across organismal clusters. Data points represent individual organisms.

**Supplementary Figure 4.**
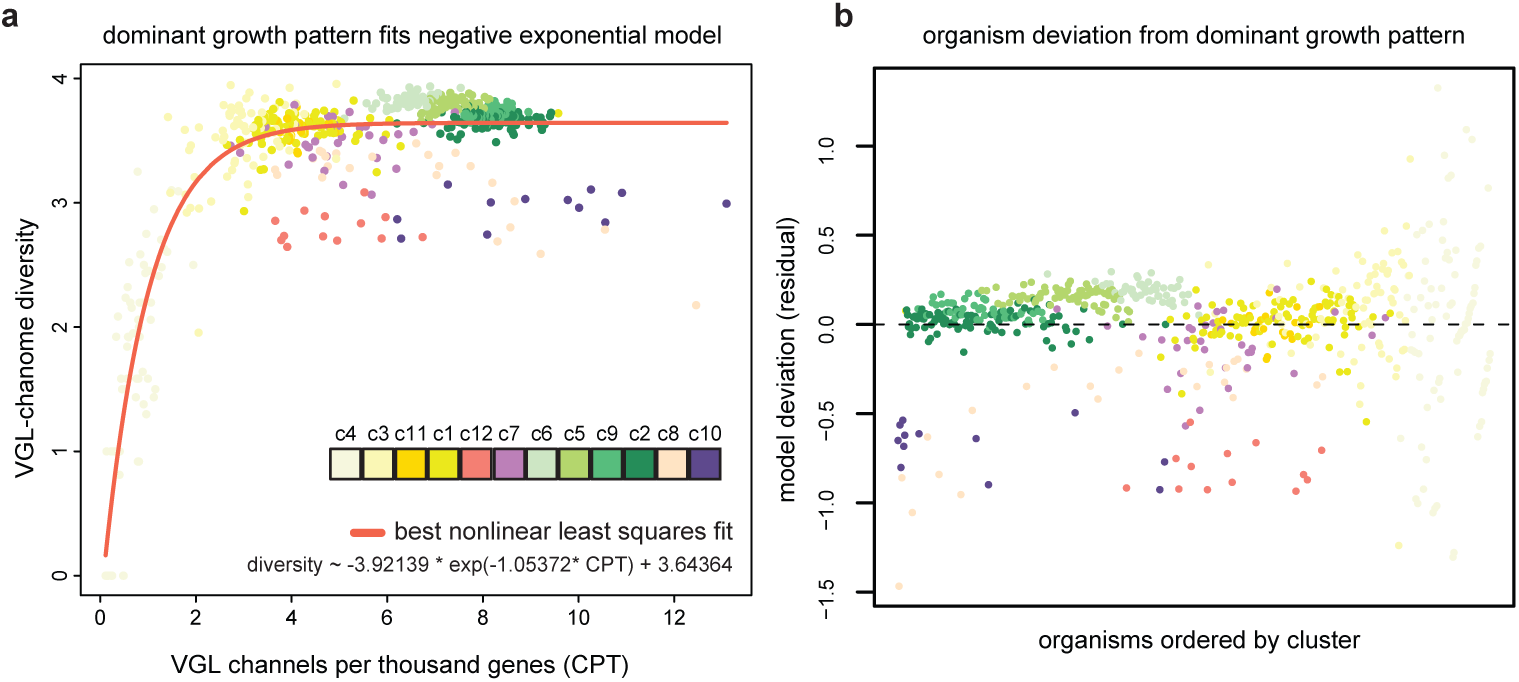
Modeling the relationship between VGL-chanome growth and diversity. **a)** Relationship between chanome size (CPT) (x-axis) and diversity (y-axis) highlighting organismal clusters (color-coded), and best-fit line of negative exponential model (tomato). Data points represent individual organisms (n=623). VGL-chanome diversity is measured using Shannon’s entropy. **b)** Deviation from predictions from the model in (**a**). Data points represent model residual values of each individual organism color-coded by organismal cluster.

**Supplementary Figure 5.**
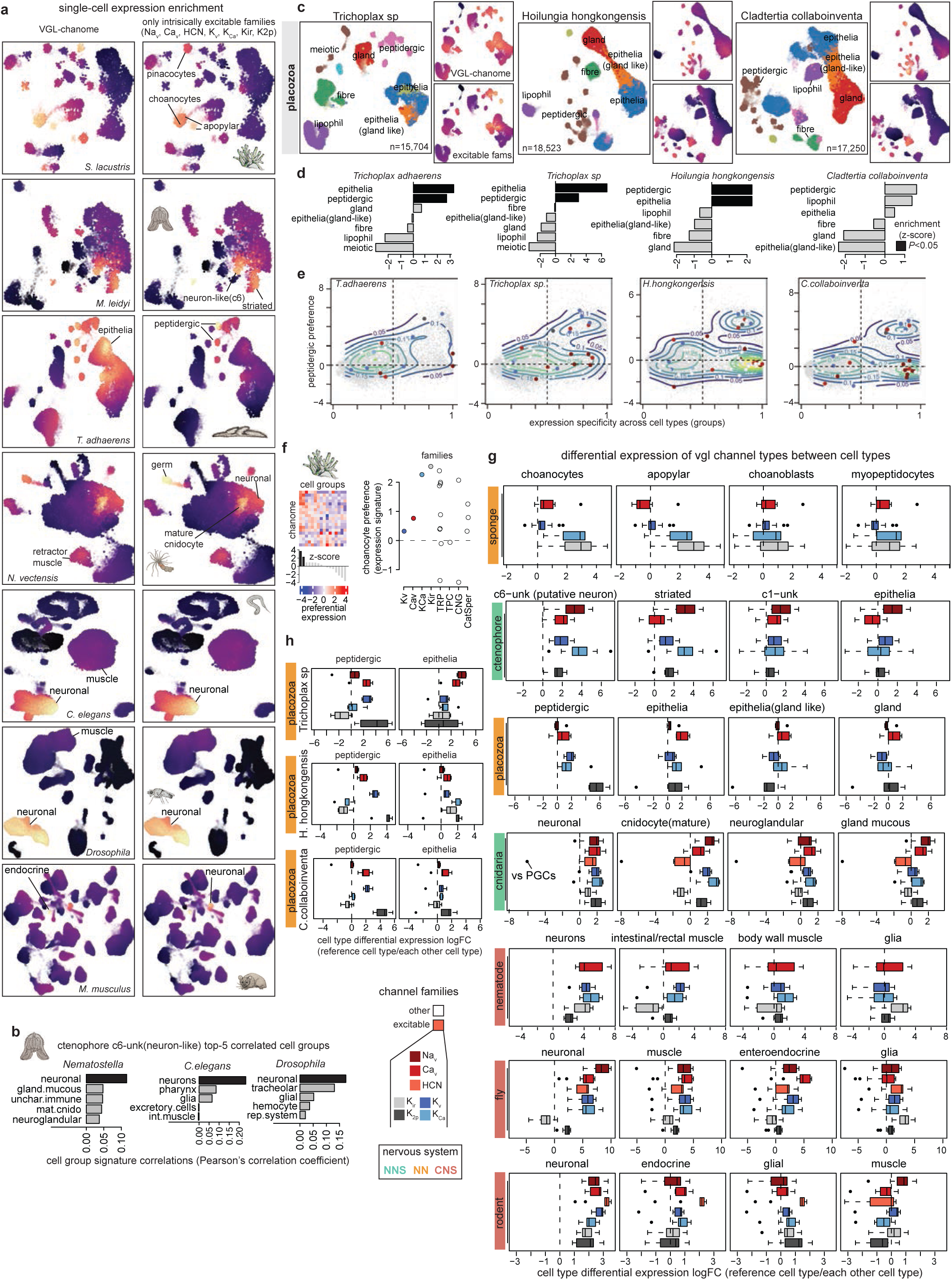
VGL-chanome preferential expression. **a)** 2-dimensional representations of single-cell transcriptomic atlases highlighting VGL-chanome single-cell expression enrichment scores. Data points represent individual cells. Data values represent enrichment of average expression values relative to random expectation. **b)** Comparison of the cell group c6-unk(neuron-like) from ctenophore versus reference transcriptomic signatures. Data values represent correlation coefficients of whole-transcriptome preferential signatures of orthologous genes. Reproducible top association highlighted in black. **c)** 2-dimensional representation of placozoan single-cell transcriptomic atlases. Data points represent individual cells labeled by major cell groups. 2D coordinates were inferred using the UMAP algorithm. Inserts highlight VGL-chanome enrichment scores. **d)** Preferential expression of VGL-chanome genes. Data values represent z-scores measuring the degree of deviation from random expectation of the average preferential expression values of VGL-chanome genes. Random expectation is estimated based on equally-sized random gene samples (n=10,000 samples with substitution). Unexpectedly high deviations are highlighted in black. *P* values were calculated based on gaussian null z-score distribution. **e)** Preference-specificity landscapes representing the relationship between the expression signature of placozoan peptidergic cells (y-axis) and gene expression specificity (x-axis). Data points represent individual genes. VGL-chanome genes are color-coded by channel family, and all genome background genes are shown in grey. Contour lines delineate areas of high density-based kernel density estimations over VGL-chanome genes. **f)** Preferential expression patterns of the VGL-chanome of *Spongilla lacustris* (*Porifera*). Data values show expression signatures across cell groups (columns) (left heatmap). Black bars represent average values over columns. Data points represent individual VGL-chanome genes (right points). **g)** Family-wise differential expression analysis comparing a reference excitable cell group (colum) versus each of the other cell groups within the organism. Data points represent pairwise comparison of the reference cell group versus another cell group. **h)** Same as (g).

**Supplementary Figure 6.**
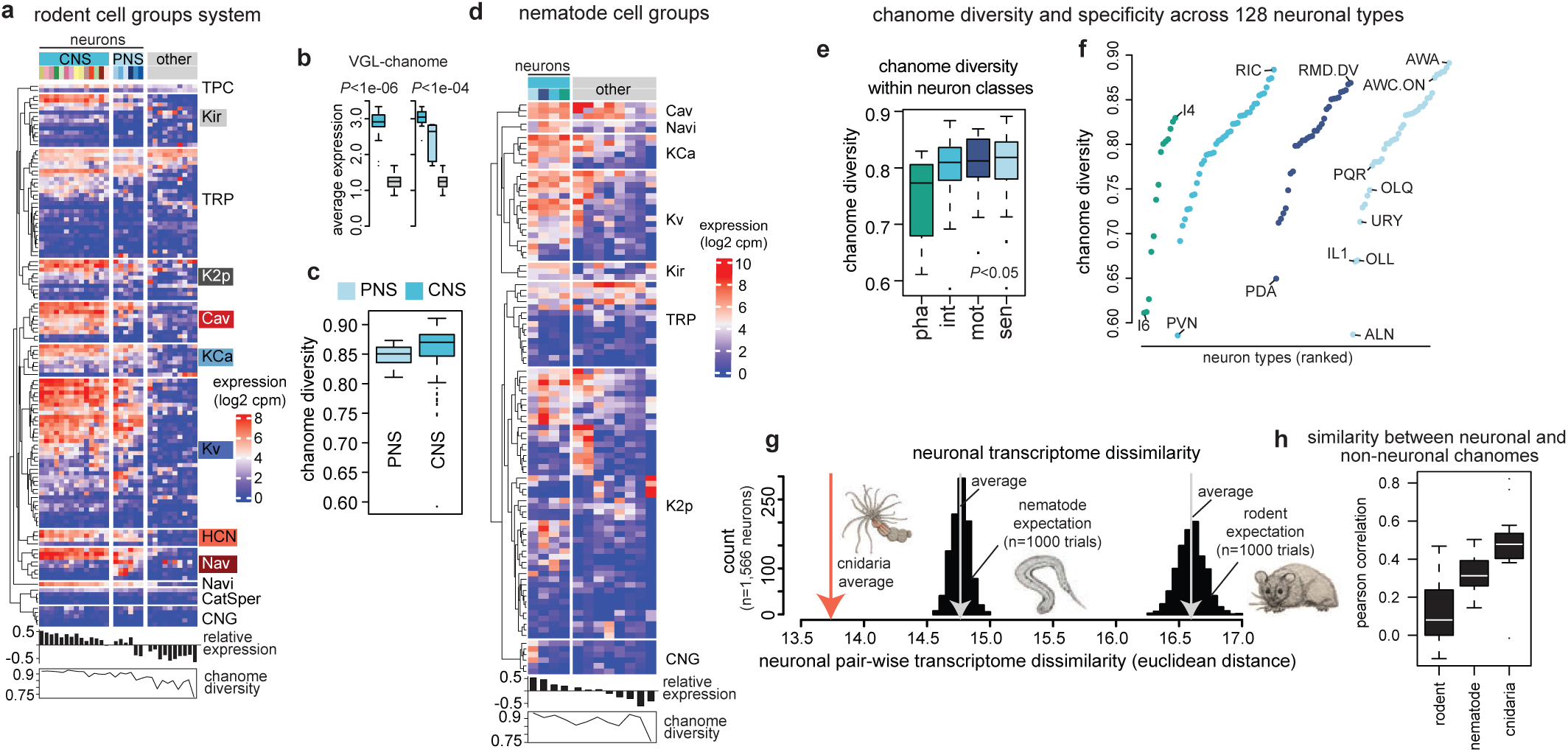
Neuronal VGL-chanome diversity. **a)** Pseudobulk VGL-chanome expression profiles of 20 neuronal taxonomic groups and 10 non neuronal cell groups of the rodent *M. musculus*. Groups corresponding to neural cells, CNS, and PNS are labeled. **b)** Average expression of VGL-chanome genes in neurons (CNS, PNS) versus non neuron groups. Data points represent individual neuron types. *P* values were calculated using Wilcoxon two-sided signed-rank tests. **c)** VGL-chanome transcriptional diversity across rodent neuron classes. Data points represent individual neuron types. Diversity is measured per neuron type using Shannon’s entropy over VGL-chanome expression profiles. **d)** Pseudobulk VGL-chanome expression profiles of 4 neuronal functional groups and 8 non neuronal cell groups of the nematode *C. elegans*. In **(a)** and **(d)** bottom black bars represent average relative expression values and bottom, line plots VGL-chanome transcriptional diversity values. **e-f**) VGL-chanome transcriptional diversity across nematode neuron functional classes **(e)** and individual neuron types **(f)**. Data points represent individual neuron types. **g**) Transcriptional dissimilarity between pairs of cnidarian, nematode, or rodent neurons. Data values represent the average pairwise Euclidean distance of 1,566 neurons (x-axis). Observed (tomato) or expected (grey) values based on random resampling are indicated with down arrows. Transcriptional dissimilarity was measured over the whole transcriptome represented in lower-dimensions (PCA, 50 PCs) using the Euclidean distance. **h**) Correlation between neuronal and nonneuronal chanomes in cnidarian, nematode, or rodent. Data values represent Pearson correlation coefficients. Correlations were measured separately per organism using pseudobulk VGL-chanome profiles.

**Supplementary Figure 7.**
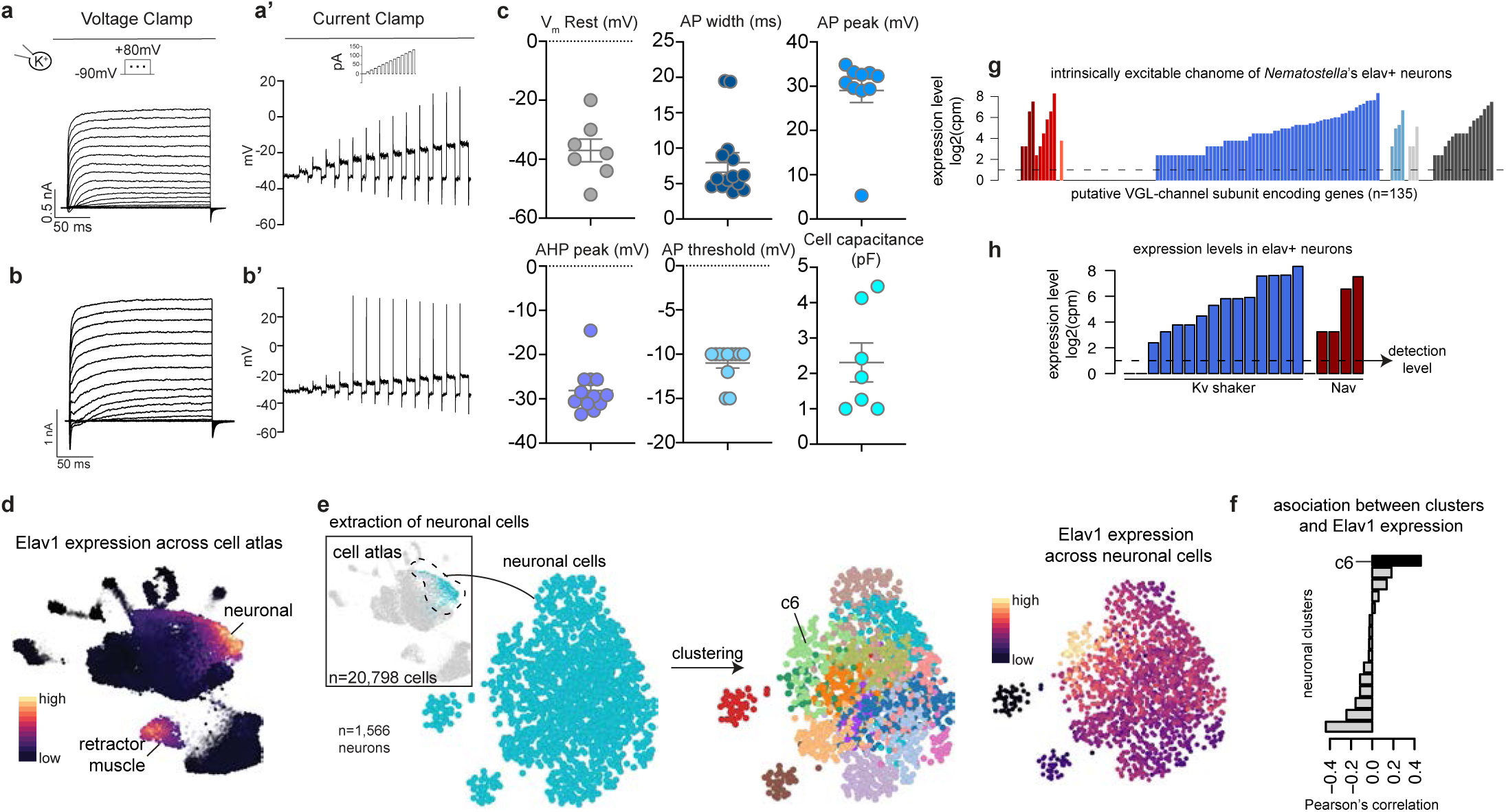
**a)** Voltage steps in Cell #1 showing current traces with **a’)** corresponding current clamp traces recording graded potentials. **b)** Voltage steps in Cell #2 showing current traces with **b’)** corresponding current clamp traces recording action potentials. **c)** quantification of action potential and cellular characteristics showing: Resting membrane potential (Vm): mean -37mV (+/- 3,861 mV, SEM), Spikewidth: mean 7,971 ms (+/- 1,387 ms, SEM), AP peak: mean 29,01 mV ( +/- 2,7 mV SEM), afterhyperpolarization (AHP): mean -28,13 mV (+/- 1,456 mV SEM), AP threshold: mean -11 mV (+/- 0.564 mV SEM), Cell capacitance: mean 2,3 pF (+/- 0,549 pF). **d)** Two-dimensional representation of single-cell transcriptomic data from Nematostella’s single cell atlas highlighting *elav1* expression values**. e)** *In silico* cell-sorting strategy to identify and extract *elav*-positive neuron in Nematostellas’ single-cell atlas. **f)** Correlation between *elav1* expression values and cluster memberships from neuronal subclustering analysis. The black bar highlights the highest correlating cluster. **g**) Expression levels in *elav1* neurons of VGL-channel subunit encoding genes. Colors represent channel families. Horizontal line represents log2(cpm)=1 detection level. **h**) Expression levels in *elav1* neurons of genes encoding K_v_-Shaker or Na_v_ subunits.

## Methods

### Datasources

#### Genomes

Reference genome annotations and protein sequences were compiled from Ensembl genome resources (https://ensemblgenomes.org/), including the specialized databases EnsemblProtists (https://protists.ensembl.org/), EnsemblMetazoa (https://metazoa.ensembl.org/), and Ensembl (vertebrates) (https://www.ensembl.org/). Only genes annotated as protein-coding were considered. The longest annotated protein sequence was used as representative protein for genes annotated with multiple transcripts/proteins. Protein sequences corresponding to gene models for organisms not present in Ensembl references and published as part of the single-cell transcriptomic studies of interest (see below) were obtained directly from the data reported in the original publication. Gene annotation and protein sources are summarized in (**Supplementary Table 1**). A total of 17,385,692 peptide sequences were downloaded. After selecting one representative sequence per annotated protein-coding gene per species, a subset of 10,849,326 unique sequences were analyzed: an average of 17,527.18 per genome (minimum 3008, maximum 49300) with an average length of 560.80 amino acids.

#### Mammalian VGL-chanome reference

A total of 145 human genes encoding voltage-gated ion channels according to The Guide to PHARMACOLOGY database of The International Union of Basic and Clinical Pharmacology (IUPHAR) / British Pharmacological Society (BPS) (version 2022.4) were obtained from https://www.guidetopharmacology.org/download.jsp. Of the initial 145 genes, the 140 voltage-gated ion channels encoding genes annotated as protein-coding in GENECODE Release 42 (GRCh38.p13), having uniquely mapped HGNC symbols and ensembl gene ids, and belonging to the families Na_v_, Ca_v_, HCN, K_v_, K_Ca_, Kir, K2p, TRP, TPC, CNG, and CatSper were used for all analyses. We refer to the resulting gene set as the reference human VGL-chanome. It includes 140 ion channel genes, classified into 11 families and 19 subfamilies (**Supplementary Table S2**).

#### Single-cell transcriptomic atlases

Whole-body single-cell RNA sequencing data of *Spongilla lacustris* (*Porifera*, sponge) was obtained from^67^. Whole-body single-cell RNA sequencing data of *Mnemiopsis leidyi* (*Ctenophora*, comb jelly) was obtained from^41^. Whole-body single-cell RNA sequencing data of placozoans (*Trichoplax adhaerens*, *Hoilungia hongkongensis*, *Trichoplax sp*., and *Cladtertia collaboinventa*) was obtained from^42^. Whole-body single-cell RNA sequencing data of *Nematostella vectensis* (*Cnidaria*, starlet sea anemone) was obtained from^46^. Whole-body single-cell RNA sequencing data of *Caenorhabditis elegans* at the L2 stage (*Nematoda*, worm) was obtained from^85^. Whole-body single-cell RNA sequencing data of *Drosophila melanogaster* (*Arthropoda*, fly) was obtained from^86^. Multi-tissue single-cell RNA sequencing data of *Mus musculus* (Mammal, mouse) was obtained from^87^. Multi-tissue single-cell RNA sequencing data of *Homo sapiens* (Mammal, human) was obtained from^88^. Single-cell RNA sequencing data of the nervous system of mouse was obtained from^44^ and of *C. elegans* from^45^. All single-cell transcriptomic data used are summarized in **Supplementary material.**

#### VGL-subfamily specific HMMs

Following (Yu and Catterall 2004), HMM models to identify putative members of the voltage-gated ion channel superfamily were based on amino acid sequences of minimal pore regions bounded by the transmembrane segments M1 or S5 and M2 or S6 of mammalian reference channel members. Models were built for 18 VGL-subfamilies (**Supplementary Table S2**). For each subfamily a HMM profile was built based on selected phylogenetic informative regions from a multiple sequence alignment. Clustal Omega (clustalo)^89^ multiple sequence alignments were made of human protein sequences of 9 Na_v_ channels, 10 Ca_v_ channels (4 Ca_v_1, 3 Ca_v_2, 3 Ca_v_3), 40 K_v_ channels (32 K_v_1-9, 8 K_v_10-12), 4 HCN channels, 15 K_ir_ channels, 8 K_Ca_ channels, 15 K_2p_ channels, 6 CNG channels, and 26 TRP channels (6 TRPC, 8 TRPM, 3 TRPML, 3 TRPP, 6 TRPV). Phylogenetic informative regions were selected using Block Mapping and Gathering with Entropy (BMGE)^90^. Hidden markov model building and search was performed using HHMER v3.4 (http://hmmer.org/). The function *hmmbuild* was used to build HMMs from multiple sequence alignment files and the function *hmmsearch* to map query HMMs to target reference proteomes.

#### Ion channel search, annotation, and filtering

To identify and score putative channel proteins in reference proteomes, each VGL-subfamily HMM model (n=18) was used as a query for searching against target species proteome sequence databases. HMM model to database searches were performed using the *hmmsearch* program of the HMMER software package for sequence analysis v3.4 (http://hmmer.org/). Each VGL-channel candidate was annotated with the best and second-best VGL-subfamily matches, best human protein hit, matches to PFAM motifs *Ion_trans* and *Ion_trans_2* motifs from Pfam (https://www.ebi.ac.uk/interpro/download/Pfam/), and number of predicted transmembrane domains (TMRs). Best human protein hits were determined by querying each putative channel sequence against the complete set of human protein-coding genes using the HMMER function *phmmer*. To minimize false positives due to partial sequences or wrong gene model predictions, only those candidates with a sequence length of at least 100 residues, a human VGL-channel as best hit, and/or a Pfam ion channel domain, a significant HMM score (HMM E-value < 0.01), and at least one predicted transmembrane domain were included in subsequent analyses. Each candidate was assigned to the best matching VGL-(sub)family for classification purposes. Candidate channels with a best model match with Ca_v_ or Na_v_ families but a best human protein match to the non-selective sodium leak channel NALCN we annotated with the Na_vi_ subfamily.

#### Membrane topology prediction

Predictions of membrane topology were performed by the deep learning protein language model-based algorithm DeepTMHMM v1.0.24^35^ using the protein sequences of all VGL- channel candidates as input. DeepTMHMM predicts one of three states for each protein residue (intercellular, extracellular, or transmembrane), along with the total number of transmembrane domains (TMR).

#### VGL-chanome profiles

A channel per thousand genes (CPT) score estimating the genome fraction encoding any VGL-channel was computed for each organism by counting the total number of VGL-channels, dividing this number by the total number of protein-coding genes in the organism, and multiplying the resulting ratio by 1000. Similar scores were computed for each VGL-channel family by performing the same computation but considering only the count of members assigned to the family -- for example, counting the number of Na_v_ encoding candidates, dividing by the total number of protein coding genes, and multiplying by 1000 results in a Na_v_PT score. Each organism was represented by a VGL-chanome profile defined as the 19-dimensional vector concatenating the normalized family scores of the 19 VGL-subfamilies (18 HMM models + Na_vi_).

#### VGL-chanome size groups

Organisms were classified in VGL-chanome investment groups by fitting a Gaussian finite mixture model to the CPT scores using the R package *mclust*. The best fit based on a Bayesian information criterion BIC -2791.472 corresponded to 4 groups.

#### Overrepresentation analysis

Statistical overrepresentation of discrete labels (e.g. taxa, VGL-chanome investment group, organismal cluster, or nervous system trait) within groups of organisms were estimated using binomial tests in all cases.

#### Organismal network construction, clustering, and visualization

Organisms were connected to each other using a weighted k-nearest neighbor (KNN) graph algorithm based on the Euclidean distance of VGL-subfamilies profiles. Each node of this graph was projected to a 2-dimensional coordinate space using the Uniform Manifold Approximation and Projection (UMAP) algorithm^91^. Both graph construction and layout estimation were performed as implemented in the *umap* R package with default parameter values. Organisms were clustered into groups by finding graph communities via random walks^92^ as implemented in the walktrap algorithm of the *igraph* R package^93^.

#### VGL-subfamily clustering relevance

VGL-subfamily relevance for organismal cluster discrimination was estimated by training a machine learning classifier to predict cluster labels based on subfamily profiles and estimate the relevance of each subfamily for the prediction task. Models were trained to do multiclass classification using the multi:softprob objective and the multiclass logloss function (*mlogloss*) as implemented in the R package *xgboost*^94^. VGL-subfamily relevance was estimated based on the relative contribution of a subfamily to the model predictions as measured by the gain metris as implemented by the *xgb.importance* function of the same package.

#### VGL-chanome diversity

VGL-chanome compositional diversity was estimated at the genomic level per organism by measuring the average level of uncertainty over VGL-subfamily size distributions using Shannon’s entropy (*H*)^95^. With 19 VGL-subfamilies, H will vary from zero when only one VGL-subfamily is present in the genome to log2(19)∼4.25 when all subfamilies are present with the same frequency. Transcriptional diversity was similarly measured by computing entropy across normalized expression profiles.

#### Modeling VGL-chanome growth and diversity

VGL-chanome diversity was modeled as a function of VGL- chanome size measured as CPT by fitting a negative exponential curve using for parameter inference the nonlinear least squares method as implemented in the *nls* function of the R *stats* package.

#### Single-cell transcriptomic data analysis

All dimensionality reduction, clustering analyses, and visualization steps were performed using the computational framework ACTIONet reported in ^96^ and available at (https://github.com/shmohammadi86/ACTIONet/tree/R-release). Each cellular landscape used for visualization and clustering was generated using the following steps. Briefly, a single value decomposition (SVD) was used to produce a low-rank approximation of the normalized count matrix. This reduced data representation is subsequently decomposed multiple times in parallel via archetypal analysis to define a multiresolution and low-dimensional cell state representation for each individual cell. This final representation is used to build a cell network or manifold capturing relationships in transcriptomic state similarity at single-cell resolution. The network is projected in 2D and 3D coordinates by the application of the UMAP algorithm. Groups of transcriptionally similar cells are obtained using the Leiden clustering algorithm. All these steps are implemented sequentially in the function *runACTIONet*. Default parameters were used in all cases. For re-analysis of *Nematostella* atlas data, the initial reduction step included a batch correction accounting for the origin of dissected tissue in adult organisms. Batch correction was implemented using ACTIONet’s function *reduce.and.batch.correct.ace.Harmony*. To score single cells using single-cell expression enrichment of VGL- chanome genes (**Supplementary Fig. S5a**), a network diffusion algorithm was first used over the corresponding cell landscape network to smooth genewise sparse expression values. These values were then averaged and compared with similar values of randomly sampled genes (n=100 replicates) to obtain a normalized enrichment z-score. Network diffusion is implemented in the ACTIONet’s *networkDiffusion* function.

#### Pseudo-bulk transcriptomic analysis

A transcriptional profile and a transcriptional signature were defined for each cell group, cell type, neuron type, or cluster. The normalized sum of total read counts per gene over cells belonging to a group was used to quantify transcriptional profiles. Read counts were normalized to counts per million (CPM) and log-transformed in base 2. A signature was defined by the normalized total sum of the pairwise expression fold-change of a given profile relative to each of the other profiles. A large positive (negative) signature value indicates increased (decreased) expression in that group over all others, with the magnitude indicating average fold-increase. Genes with high signature values for a given group represent preferentially expressed genes and tend to correspond to well-known cell type markers. VGL-chanome expression enrichment (**Fig. 5b**) was assessed by contrasting observed average transcriptional signature values with a null distribution estimated by randomly sampling same-sized gene sets (10,000 random trials with substitution) and quantifying the deviation from mean expectation with a z-score. Transcriptional signatures were used as measures of cell group preference. To measure gene expression specificity, a gene specificity score was computed using the *Tau* method as described in^97^. Briefly, *Tau* summarizes in a single number whether a gene tends to be cell group specific (close to 1) or ubiquitously expressed (close to 0) across conditions (cell groups).

#### Orthology inference

Orthologous groups used for comparative single-cell transcriptomic analysis were inferred using OrthoFinder version 2.5.5^98^. Orthologous groups with unique genes for the pair of species to be compared when considered when comparing cell group signatures.

### Animals and Cells

*Nematostella vectensis* (*Elav1::mOrange*) were a gift from the Rentzsch lab (University of Bergen). Animals were maintained according to Stefanik, Friedman and Finnerty (Stefanik et al., 2013) and communicated instructions from the Rentzsch lab: in brief the sea anemones were kept in glass Pyrex dishes filled with ⅓ NSW (Seawater obtained at Trondheim Biological Station from 100m depth and sterile filtered) at room temperature in a 12/12 light dark cycle. The anemones were fed two to three days a week with freshly hatched brine shrimp nauplii (Artemia sp.). Cells were isolated from transgenic animals as follows: animals were allowed to relax in ⅓ NSW. Upon relaxation Nematostella anesthesia solution (containing (in mM):140 NaCl, 3.3 KCl, 3.3 HEPES, 3.3 Glucose, 40 MgCl2, pH 7.6) was added and tentacles were dissected from the animals using a syringe needle. The tentacles were transferred to TrypLE Select 10x (Gibco, A12177-01) for 30 min @ 32 C with regular agitation. Cells were then dissociated in divalent free solution (containing (in mM):140 NaCl, 3.3 KCl, 3.3 HEPES, 3.3 Glucose, pH 7.6). Dissociated cells were kept in a holding solution supplemented with glucose to an osmolarity of 305-315 mOsm/Kg on ice until use.

### Electrophysiology

Patch clamp recordings were carried out at room temperature using a MultiClamp 700B amplifier (Axon Instruments) and digitized using a Digidata 1550B (Axon Instruments) interface and pClamp software (Axon Instruments). Whole-cell recording data were filtered at 1kHz sampled at 10 kHz. Voltage-gated currents were leak-subtracted online using a p/4 protocol. For Nematostella cell recordings, borosilicate glass pipettes were polished to 9-11MΩ. The standard extracellular solution contained (in mM): 140 NaCl, 3.3 KCl, 3.3 HEPES, 3.3 Glucose, 2 CaCl2, 0.5 MgCl2, pH 7.6. The intracellular solution used to measure K+ currents and membrane voltage, contained (mM): 150 K+ gluconate, 3.3 HEPES, 10 sucrose, 1.33 MgCl2, 3.3 KEGTA, pH 7.6, in some experiments ATP was added to prevent inward current rundown.

## Resource availability

### Lead contact

Further requests for resources and reagents, data, or code should be directed to and will be fulfilled by the lead contact, Jose Davila Velderrain

### Materials availability

This study did not create new unique reagents.

### Data and code availability

VGL-channel annotations, protein sequences, topology predictions, summary information of transcriptomic data sources, processed data, and annotation tables will be available at Mendeley Data upon publication. Any additional information required to reanalyze the data is available upon request.

## Acknowledgements

We thank Fabain Rentzsch (UiB) for gifting the *Elav1::mOrange* animals. This research was supported by the ERC StG “EnvIronchannel” (101076516) granted to L.v.G. and by the Human Technopole Foundation supporting J.D.V.

## Author contributions

J.D.V performed computational analysis, L.v.G performed electrophysiological and imaging experiments. All authors conceptualized the study, designed and created the figures, and wrote the manuscript.

